# Structural insights into the molecular mechanisms of the *Mycobacterium* evolvability factor Mfd

**DOI:** 10.1101/728246

**Authors:** Sivasankar Putta, Swayam Prabha, Vinayak Bhat, Gavin C. Fox, Martin A. Walsh, Desirazu N. Rao, Valakunja Nagaraja, Ramanathan Natesh

**Affiliations:** School of Biology, Indian Institute of Science Education and Research Thiruvananthapuram, Thiruvananthapuram 695551, India; Department of Biochemistry, Indian Institute of Science, Bangalore 560012, India; Department of Microbiology and Cell Biology, Indian Institute of Science, Bangalore 560012, India; Synchrotron SOLEIL, L’Orme des Merisieris Saint-Aubin BP48, 91192 Gif-sur-Yvette, France; Diamond Light Source, Didcot OX11 0DE & Research Complex at Harwell, Didcot OX11 0FA, UK; Jawaharlal Nehru Centre for Advanced Scientific Research, Bangalore 560064, India

**Keywords:** crystal structures, negative stain and cryo-EM structure, DNA damage, Transcription-Coupled Repair, Transcription Elongation Complex, Stalled RNAP, Electrophoretic Mobility Shift Assay (EMSA), MtbMfd oligomer, *M. tuberculosis* Mfd and *M. smegmatis* Mfd structures

## Abstract

Mfd is a highly conserved ATP dependent DNA translocase that mediates the role of Transcription-Coupled-DNA-Repair(TCR) in bacteria. The molecular mechanisms and conformational remodelling that occurs in Mfd upon ATP binding, hydrolysis, and DNA translocation are poorly defined. Here we report a series of crystal and electron microscopy(EM) structures of Mfd from *Mycobacterium tuberculosis* (MtbMfd) and *Mycobacterium smegmatis* Mfd, solved in both the apo and nucleotide-bound states. The structures reveal the mechanism of nucleotide-binding, which lead to the remodeling of the Walker A motif at the catalytic pocket, inducing a flip-flop action of the hinge and flexible linker regions. Specifically, nucleotide binding unlocks the Translocation in RecG motif of the D6-domain to induce a ratchet-like motion. Functional studies of MtbMfd-RNAP complexes show that MtbMfd rescues stalled Transcription Elongation Complexes. We also report negative-stain and cryo-EM single particle reconstructions of MtbMfd higher order oligomer, that reveal an unexpected dodecameric assembly state. Given that Mfd accelerates the evolution of antimicrobial resistance(AMR), our results establish a framework for the design of new “anti-evolution” therapeutics to counter AMR.

## INTRODUCTION

TCR is a DNA damage repair mechanism found in all cells. In humans TCR is implicated in Cockayne Syndrome, which is manifested by accelerated aging and is fatal in early life (Cleaver, Lam et al., 2009). In bacteria, a product of the *Mfd* gene called transcription-repair coupling factor (TRCF) or Mfd, mediates TCR. Mfd interacts with stalled RNA polymerase (RNAP) at the site of DNA damage and helps to nudge the RNAP back into position to resume productive elongation. If this fails, it then mediates the removal of the stalled RNAP and recruits the DNA repair proteins to the site of damage. Thus, Mfd both clears steric hindrance from the site of damage, and if necessary, loads the UvrABC repair protein complex, resulting in repair of the damaged strand (Deaconescu, Chambers et al., 2006, Le, Yang et al., 2017, Park, Marr et al., 2002, Selby & Sancar, 1993, Svejstrup, 2002, Trautinger, Jaktaji et al., 2005). It has recently been shown that Mfd employs a ‘‘release and catch-up’’ mechanism to rescue stalled or backtracked Transcription Elongation Complex (TEC) (Le et al., 2017). Mfd catalyses two irreversible ATP-dependent transitions that lead to the removal of stalled RNAP from the DNA damage site (Fan, Leroux-Coyau et al., 2016, Graves, Duboc et al., 2015, Howan, Smith et al., 2012). The eXcision Repair-sequencing method revealed that Mfd mediates genome-wide TCR (Adebali, Chiou et al., 2017). Mfd is a highly conserved central player in bacterial TCR with putative functions in processes outside of DNA damage, such as to resolve conflicts between DNA replication and transcription(Trautinger et al., 2005), carbon catabolite repression(Zalieckas, Wray et al., 1998) etc. Our current understanding on structure, function and mechanism of Mfd mediated TCR is based largely on studies with *E. coli* Mfd (EcMfd). EcMfd is a monomeric protein and has been extensively investigated to understand the Mfd reaction mechanism in TCR (Deaconescu et al., 2006, Fan et al., 2016, Graves et al., 2015, Howan et al., 2012, Park et al., 2002, Selby, 2017, Selby & Sancar, 1993, Svejstrup, 2002). *In vivo* studies have shown that mutations in *E. coli mfd* leads to modest sensitivity to UV in wild-type cells whereas it confers high sensitivity in *recA ^−^* cells (Selby & Sancar, 1993). Recent studies have shown that Mfd also acts as an “evolvability factor” promoting antibiotic resistance in diverse bacterial species, including *M. tuberculosis* (Ragheb, Thomason et al., 2019). The emergence of multi drug resistant strains of *M. tuberculosis* and other pathogenic microorganisms is an increasingly urgent threat to human health worldwide. In this light, our investigation provides new structural, functional and detailed mechanistic information on Mycobacterial Mfd, which can provide a framework for targeted design of new classes of drugs that inhibit AMR.

In this study, we present the crystal structures of *Mycobacterium tuberculosis* Mfd (MtbMfd) and *Mycobacterium smegmatis* Mfd(MsMfd) in apo and nucleotide-bound forms. The structures reveal in atomic detail the intricate and distinctive dynamics of Mfd upon nucleotide binding/hydrolysis, mechanistic insight into a sequence of domains associations/dissociations, plausible mechanism for DNA translocation and a previously undiscovered role of D3 domain in nucleotide binding. *In vitro* functional studies suggest that MtbMfd displacement activity is optimal in the presence of mycobacterial RNAP as opposed to EcRNAP, which further support the hypothesis that Mfd and RNAP have phyla-specific interactions. Negative stain and cryo electron microscopy studies of Mtb Mfd oligomer demonstrate the existence of a hitherto unknown dodecameric form of MtbMfd.

## RESULTS

### Crystal structures of *Mycobacterium* Mfd’s and its domain organization

Using an integrated structural biology approach, we have solved the crystal structures of the apo and nucleotide-bound forms of monomeric MsMfd (131 kDa) and MtbMfd (133 kDa) and the structure of the oligomeric form of MtbMfd by electron microscopy (EM) (Figures 1A-D, S1 and S3). Ab-initio phasing of MsMfd was carried out using selenomethionyl-substituted crystals (MsMfd^se^) and the resolution was extended to 2.75 Å by Molecular Replacement (MR) using a native data set. MtbMfd and binary complexes of MsMfd with Adenosine Diphosphate (ADP) (MsMfd^ADP^) and ATPgammaS (MsMfd^ATPgammaS^) were then solved by MR to 3.6 Å (Figure 1C), 3.5 Å (Figure 1D) and 3.3 Å resolution (not shown), respectively. The crystallographic data and refinement statistics are summarized in Table 1. The crystal structures reveal the molecular basis of nucleotide binding (Figures 2B-2D and 3B; Movies S2 and S3), and suggest a plausible DNA translocation mechanism (Figures 3C and 4E). Single particle negative stain and cryo electron microscopy (cryo-EM) and three dimensional image processing were used to reconstruct a 3D image of the oligomeric form of MtbMfd (Figures 1E and S3A-G).

**Figure 1.**
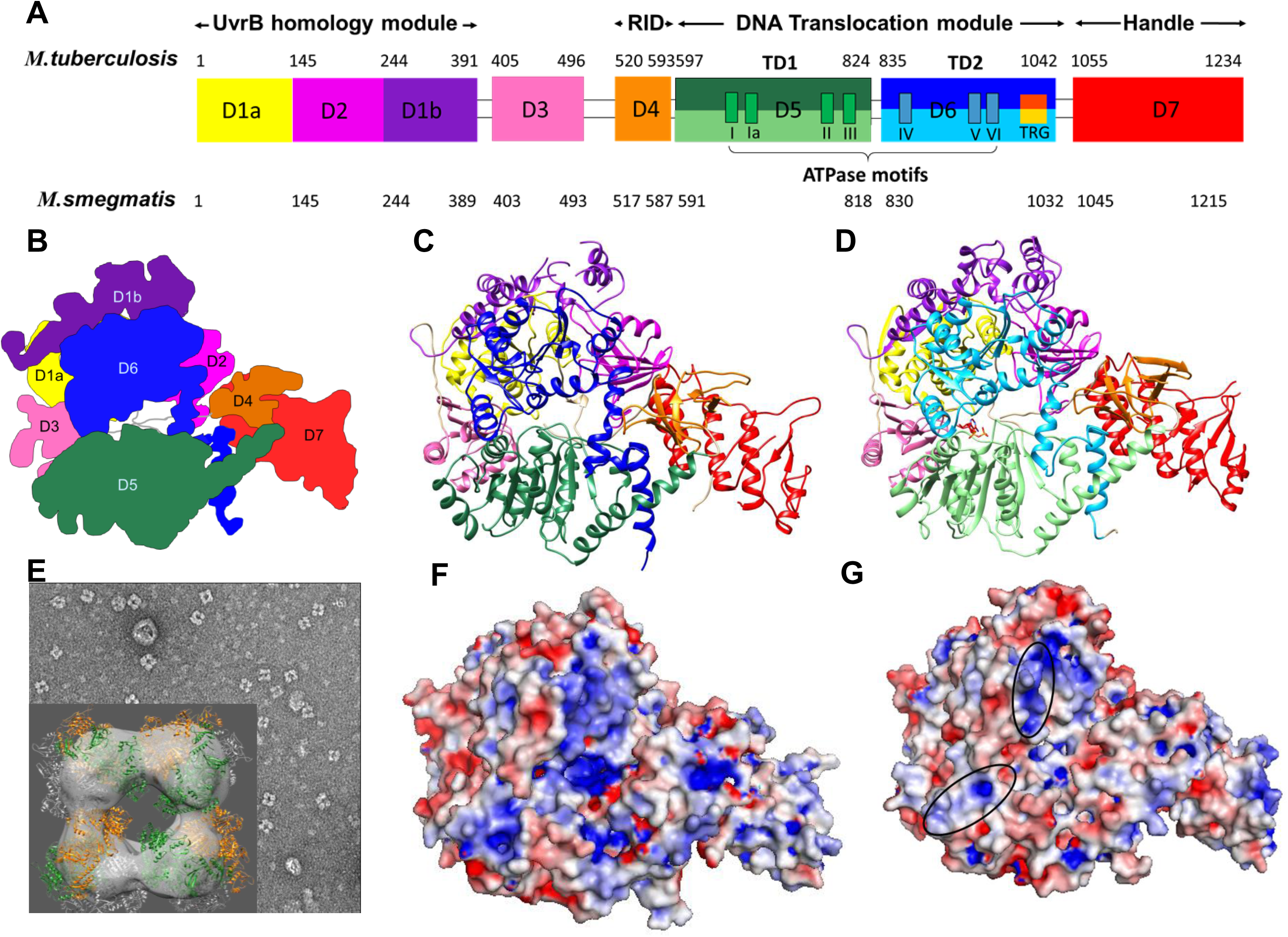
**The structures of Mfd from Mycobacterium tuberculosis and Mycobacterium smegmatis:** **(A)** Schematic diagram of the domain organization of MtbMfd and MsMfd (domain colours: D1a (yellow), D2 (magenta), D1b (purple), D3 (hotpink), D4 (orange), D5 (light green), D6 (skyblue) and D7 (red). The numbers above and below the horizontal bar represent domain ranges in MtbMfd and MsMfd. The conserved seven ATPase motifs of both TD1 and TD2(Deaconescu et al., 2006) are colored green and cornflower blue. The TRG (translocation in RecG) motif(Deaconescu et al., 2006, Mahdi et al., 2003) of the D6 domain of *M. tuberculosis* and *M. smegmatis* are orange and gold. **(B)** Schematic of the architecture of the *Mycobacterial* Mfds. **(C,D)** Ribbon models of apo MtbMfd and MsMfd^ADP^ determined at 3.6 Å and 3.5 Å resolution. The 8 domains are shown in different colours as shown in **(A)**. **(E)** Negative stain EM 3D reconstruction of the MtbMfd oligomer. **(F, G)** Electro static surface potential of MtbMfd and MsMfd. The proposed DNA interacting surface area composed of conserved basic residues is indicated by the black circle. The Mfd’s electrostatic surface potentials were calculated with the APBS plug-in in pymol (www.pymol.org), and are contoured from –5 (red) to +5 (blue) kT/q.

**Figure 2.**
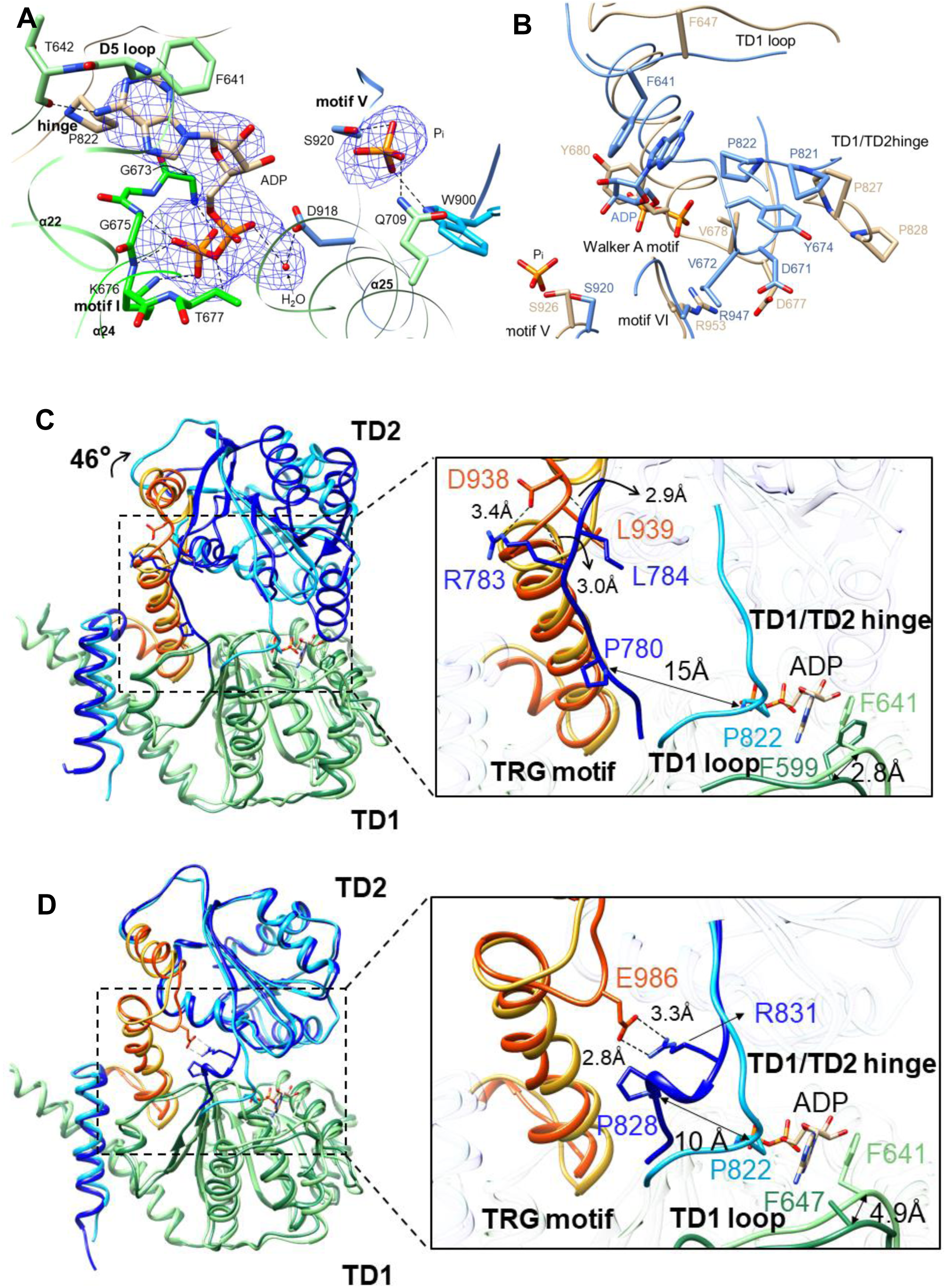
**Conformational switching in Mfd upon nucleotide binding.** Domains and motifs are coloured as in Figure 1A. **(A)** ATP binding pocket of Mfd translocation module. *2mF_o_-DF_c_*composite omit map for MsMfd bound ADP, P_i_ and Water (H_2_O) are shown as blue mesh contoured at the 1σ level (*F_o_-F_c_*map is shown in Figure S1G). Black dashed lines represent hydrogen bonds. **(B) The Walker A motif switches in response to nucleotide binding.** Superposition of MsMfd^ADP^ (cornflower blue), and apo-MtbMfd (tan) crystal structures reveal the nucleotide-induced conformation in the Walker A motif. In the absence of ADP, the Walker A motif forms a compact helical conformation wherease in the presence of ADP, it adopts an extended conformation, which results in stacking of the conserved Tyr674 in Walker A motif and Pro821 in the hinge-region. The conserved Tyr (Tyr680 in MtbMfd) in the apo MtbMfd positions into the nucleotide binding region and occludes the nucleotide binding site (See Movie S4). The MsMfd^ADP^ Arg947 of the conserved arginine finger in motif VI which senses ATP hydrolysis for Mfd translocation(Chambers et al., 2003, Deaconescu et al., 2006, Mahdi et al., 2003), lies proximally to the nucleotide binding pocket. In apo MtbMfd Arg953 (Arg947 in MsMfd^ADP^) forms a strong salt bridge with the backbone carbonyl oxygen of Asp677 (Asp671 in MsMfd^ADP^), with a Psi angle of Asp677 in a cis conformation (which we propose is due the the proximity of Arg953 as a result of TD2 crystal contact). **(C, D)** Nucleotide induced conformational changes in the translocation module of Mfd. **(C)** Superposition of the translocase module of apo EcMfd (TD1-sea green and TD2-blue) onto MsMfd^ADP^ (TD1-light green and TD2-deep sky blue), shows a rotation of the EcMfd-TD2 domain by ∼ 46° w.r.t. the TD2 domain of MsMfd^ADP^ (See also Movie S1). However, the apo MtbMfd did not show any TD2 rotation (Figure 2D), which might be due to crystal contacts at the TD2 domain (Figure S5A). Expanded view shows the conformations of the TD1 loop and hinge-regions of EcMfd and MsMfd^ADP^. The conserved Phe599 from the TD1 loop region and the Pro780 of the hinge-region are 2.8 Å and 15 Å away from the corresponding residues in MsMfd^ADP^. The hinge (blue) makes contacts (black dashed lines) with the TRG motif (orange red). **(D)** Superposition of the translocase module of apo MtbMfd (TD1-sea green and TD2-blue) on MsMfd^ADP^ (TD1-light green and TD2-deep sky blue). Right, close-up view shows conformations of the TD1 loop and the hinge region of apoMtdMfd and MsMfd^ADP^. In apo MtbMfd, the hinge (blue) makes a strong bidentate salt bridge (black dashed lines) with the TRG motif (orange red). In the case of MsMfd^ADP^the hinge region (deep sky blue) and concurrently the TD1 loop are displaced by 10 Å and 4.9 Å, respectively (11.8 Å and 7.9 Å repectively when full length Mfd structures are aligned (Figure 3B), closer towards the nucleotide binding pocket. This forces Pro822 and Phe641 to stack against ADP as described in (B), and main text. The loss of the hinge region contact with the TRG motif upon nucleotide binding therefore acts as a conformational switch.

**Figure 3.**
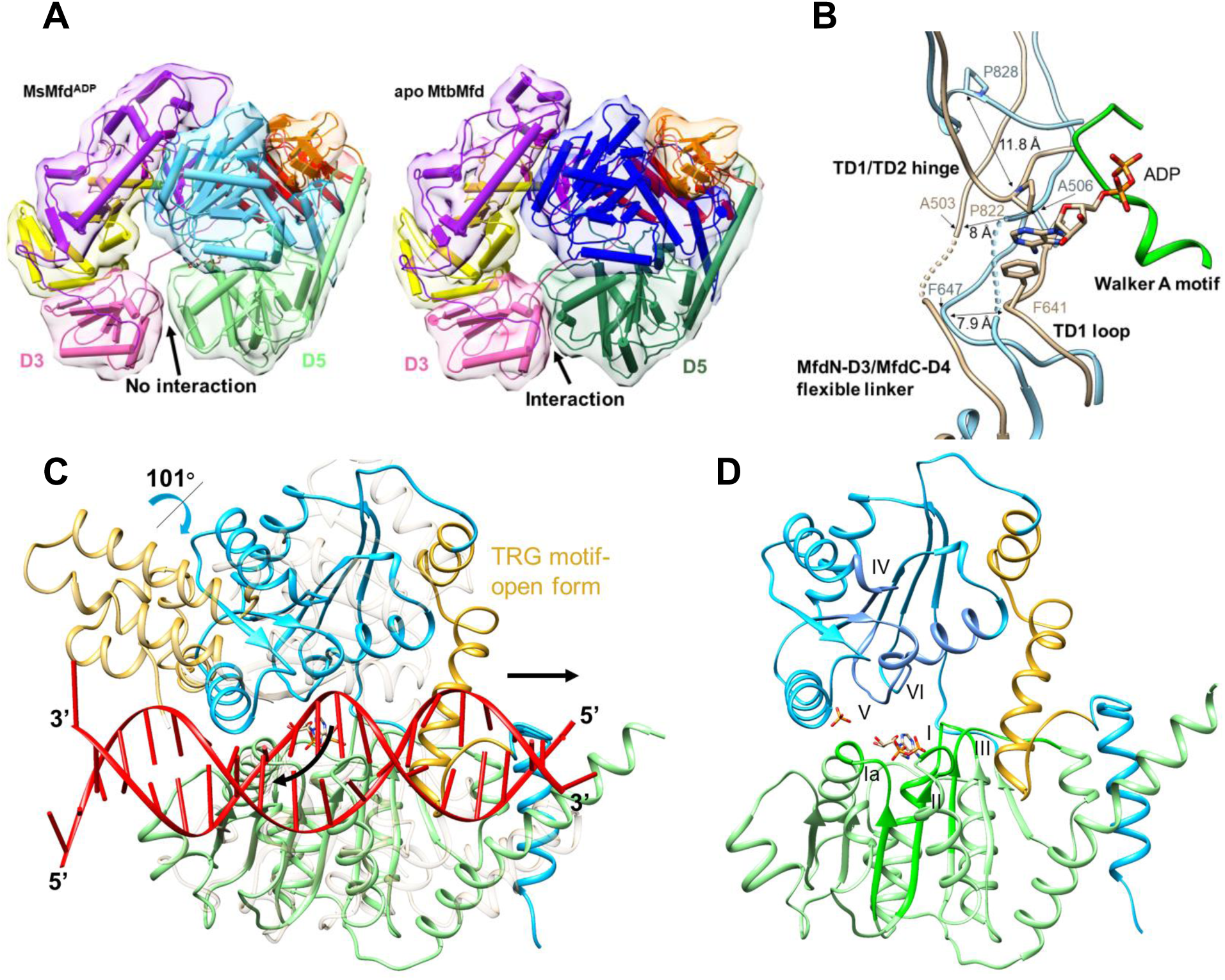
**Mechanistic insight into Mfd nucleotide binding and DNA translocation.** **(A)** Loss of interactions between D3 and D5 domains upon nucleotide binding (left panel) w.r.t. apo form (right panel, Table S1C). **(B)** Flip-Flop movement of the flexible linker and hinge-region in the ATP catalytic cycle. In MsMfd^ADP^ (tan) vs MtbMfd (light blue), displacements of the hinge by 11.8 Å and the TD1 loop by 7.9 Å towards the nucleotide, to stack Pro822 (hinge) and Phe641 (TD1 loop) against the adenine base of ADP are illustrated. Simultaneously the flexible linker is displaced away by ∼ 8 Å to accommodate the incoming hinge and TD1 loop to stack against the bound nucleotide. **(C)** Modelled MsMfd^ADP^-DNA complex. The model was generated by superposition of the SsRad54-dsDNA(Durr et al., 2005) translocase module (shown as transparent in background) onto the MsMfd^ADP^ TD1 (light green) / TD2 (deep sky blue). MsMfd^ADP^ TD2 displays a 101° rotation with respect to nucleotide-free SsRad54-dsDNA-TD2 (when TD1 domains are superposed) and is positioned closer to the bound DNA. The open TRG motif in MsMfd^ADP^ is coloured gold and the corresponding closed subdomain 2B of SsRad54 is transparent gold. Curved and straight black arrows point to the DNA transport and Mfd translocation directions, respectively. **(D)** The RecA-like translocation (motor) domains and their ATPase motifs in MsMfd^ADP^. The ATPase motifs of the TD1 (light green) and TD2 (deep sky blue) domains are shown in green and cornflower blue. The TRG motif is shown in gold. The ATPase motifs I to VI are juxtaposed with the nucleotide-binding pocket in contrast with apo-EcMfd (Figure S7B).

**Figure 4.**
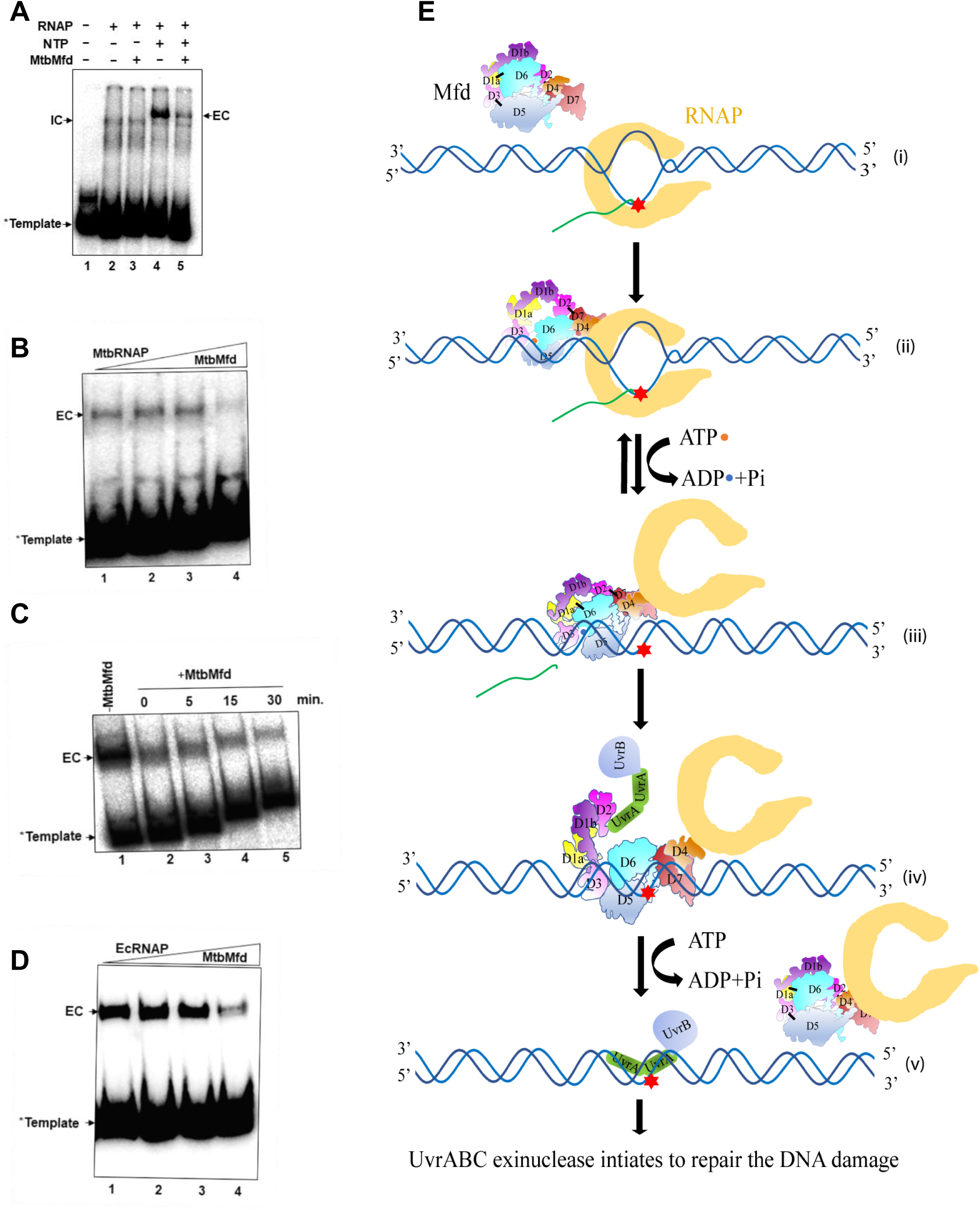
**RNAP displacement activity of MtbMfd and model for mechanism of Mfd in TCR.** **(A)** Effect of MtbMfd on Initiation Complex (IC) and Elongation Complex (EC). Lane 1, DNA template alone; Lane 2 & 3, IC in the absence and presence of MtbMfd, lane 4 & 5 EC in absence and presence of MtbMfd (* depicts the end labeled template). The loss of MsRNAP displacement in the IC is probably due to the presence of sigma factor that could hinder the binding of MtbMfd to the upstream DNA. **(B,D)** MtbMfd-mediated displacement of **(B),** MtbRNAP and **(D),** EcRNAP *in vitro*. For both **(B)**, and **(D)**, Lane 1, in each figure is without MtbMfd; Lanes 2-4 are in the presence of increasing concentrations 100, 250 and 1000 nM of MtbMfd. MtbMfd can displace the MtbRNAP more effeciently than the EcRNAP, indicating that Mfd directed RNAP interaction is phylum specific. **(C)** Time dependent displacement of MsRNAP by MtbMfd. Lane 1, the reaction was stopped at 0-time point by addition of 1 x DNA loading dye. Lane 2-4, reactions were stopped at 5, 15 and 30 min. The release of RNAP from the template DNA was scored by EMSA (4 %) at 4 °C and visualized by phosphor imaging. **(E)** Proposed mechanistic model of Mfd domains association and dissociation upon ATP binding, hydrolysis and DNA translocation in TCR. The D1:D6, D2:D7 and D3:D5 interactions (shown as thick short black line) exist in the apo form of Mfd (i). Mfd in complex with the stalled RNAP and ATP might allow the conformational changes that disrupt the D1:D6 and D3:D5 interactions (ii). In this conformational state, the D6 of Mfd may tune its position to locate the TRG motif (Pawl) on the upstream minor groove of DNA to initiate the Mfd translocase movement (in a ratchet like motion) during the ATP-ADP catalytic cycle (ii). In the ADP bound state, the D1-D6, and D2-D7 interactions remained intact (iii). ADP to ATP exchange allows to step on to the new translocated upstream DNA site. Upon signaling the damage site, Mfd recruits the UvrA2-B complex by binding to D2 which thereby unmask the D2 domain from the D7 domain(Deaconescu et al., 2006, Fan et al., 2016, Graves et al., 2015, Howan et al., 2012, Selby & Sancar, 1995) (iv). Finally, ATP hydrolyis leads to the Mfd-RNAP complex detaching from the DNA(Fan et al., 2016) (v). The UvrA2-B complex intiates the UvrABC-mediated DNA excision machinery to repair the DNA damage.

**Table 1.** Crystallographic data collection and refinement statistics

|  | MsMfd | MsMfd<sup>Se</sup> | MsMfd<sup>ADP</sup> | MsMfd<sup>ATPgammaS</sup> | MtbMfd |
| --- | --- | --- | --- | --- | --- |
| **Data collection** |  |  |  |  |  |
| Space group | P2<sub>1</sub>2<sub>1</sub>2<sub>1</sub> | P2<sub>1</sub>2<sub>1</sub>2<sub>1</sub> | P2<sub>1</sub>2<sub>1</sub>2<sub>1</sub> | P2<sub>1</sub>2<sub>1</sub>2<sub>1</sub> | P432 |
| Molecules per ASU | 2 | 2 | 2 | 2 | 1 |
| Cell dimensions |  |  |  |  |  |
| *a*, *b*, *c* (Å) | 83.9,162.0,215.4 | 82.7,158.9,207.9 | 83.7, 160.4,214.5 | 84.4,164.2, 216.2 | 224.3,224.3,224.3 |
| $\alpha$, $\beta$, $\gamma$ (°) | 90.0, 90.0, 90.0 | 90.0, 90.0, 90.0 | 90.0, 90.0, 90.0 | 90.0, 90.0, 90.0 | 90.0, 90.0, 90.0 |
| Resolution (Å) | 40.59 - 2.75 | 50.00 - 2.99 | 31.73 - 3.50 | 39.81 – 3.30 | 39.64 - 3.60 |
| *R*merge (%) | 7.6 (82.6) | 13.2 (113.7) | 20 (59.4) | 8 (50.4) | 14.3 (100.6) |
| *I*/σ (*I*) | 9.0 (1.6) | 14.4 (1.9) | 6.3 (2.4) | 12.0 (2.5) | 8.7 (2.2) |
| CC<sub>1/2</sub> | 0.99 (0.46) | 0.99 (0.76) | 0.98 (0.35) | 0.99 (0.41) | 0.99 (0.38) |
| Completeness (%) | 99.5 (99.7) | 99.0 (92.4) | 96.8 (97.1) | 81.3 (64.7) | 99.6 (99.3) |
| Redundancy | 4.5 (4.3) | 13.7 (13.64) | 5.1 (4.9) | 4.7 (3.2) | 8.7 (8.7) |
| **Refinement** |  |  |  |  |  |
| Resolution (Å) | 40.50 - 2.75 | 47.19 - 2.99 | 30.10 - 3.50 | 39.3 - 3.3 | 39.64 - 3.6 |
| No. reflections<br>(Total/test) | 76560 / 3815 | 55718 / 2787 | 35902 / 1768 | 37322 / 1800 | 22838 / 1116 |
| *R*work / *R*free (%) | 22.4 / 25.7 | 21.8 / 26.9 | 22.6 / 27.3 | 22.0 / 27.1 | 24.9/29.5 |
| **No. atoms in ASU** |  |  |  |  |  |
| Protein | 17156 | 16754 | 17167 | 17149 | 8259 |
| Ion & Ligand | 55 & 0 | 25 & 0 | 25 & 54 | 25 & 31 | 0 & 0 |
| Water | 120 | 74 | 105 | 37 | 16 |
| **Average B-factors** |  |  |  |  |  |
| Protein (Å<sup>2</sup>) | 84.6 | 80.2 | 69.6 | 63.9 | 114.5 |
| Ion & ligand (Å<sup>2</sup>) | 102.1 | 84.1 | 91.9 | 71.4 | - |
| Water (Å<sup>2</sup>) | 69.7 | 64.1 | 33.1 | 37.6 | 74.7 |
| **R.m.s. deviations** |  |  |  |  |  |
| Bond lengths (Å) | 0.003 | 0.003 | 0.002 | 0.002 | 0.002 |
| Bond angles (°) | 0.59 | 0.58 | 0.52 | 0.47 | 0.57 |
| **Ramachandran plot statistics (%)**<sup>#</sup> |  |  |  |  |  |
| Most favored region | 89.4 | 89.2 | 90.6 | 89.0 | 84.1 |
| Additionally allowed Region | 10.4 | 10.6 | 9.1 | 10.8 | 15.9 |
| Generously allowed Region | 0.2 | 0.2 | 0.3 | 0.2 | 0.0 |
| Disallowed region | 0.0 | 0.0 | 0.0 | 0.0 | 0.0 |
|  | Values in parentheses are for the highest-resolution shell.<br><sup>#</sup> Calculated for non-glycine and non-proline residues using PROCHECK |  |  |  |  |

In spite of functional similarities with to EcMfd, biochemical analysis shows that MtbMfd has distinct properties (Prabha, Rao et al., 2011). It was shown that MtbMfd exists in both monomeric and hexameric forms *in vivo* and *in vitro* (Prabha et al., 2011), while EcMfd exists only in monomeric form (Deaconescu et al., 2006). Similar to *E.coli* Mfd (Deaconescu et al., 2006), *Mycobacterial* Mfd is comprised of eight domains (Figure S2). The N-terminal region (MfdN) includes domains D1a - D3, connected by a flexible linker to the C-terminal region (MfdC) incorporating the D4 - D7 domains (Figures 1A and 1B). The N-terminal domain (NTD) (D1a, D2, and D1b) of Mfd is a structural homologue of UvrB and known for its interaction with the UvrA. The D3 domain has been proposed to be species-specific with unknown function. The RNA polymerase interaction domain (RID) (D4) plays a critical role during TCR by interacting with the β-subunit of the RNA polymerase. The D5 and D6 of C-terminal region of Mfd harbours the ATPase and TRG (translocation in RecG) motifs that are essential for the function of ATP dependent DNA translocation. The D7 domain is considered to be auto inhibitory domain in Mfd function (Deaconescu et al., 2006, Selby & Sancar, 1995).

### The conformation of the Translocation module and its dynamics upon nucleotide binding

Mfd belongs to the superfamily 2 (SF2) helicases (Selby & Sancar, 1995, Svejstrup, 2002). In SF1 and SF2 helicases, the two RecA-like domains play a vital role in the ATPase activity as well as in DNA-binding (Caruthers & McKay, 2002). Although, a crystal structure of *E. coli* Mfd alone has been determined (Deaconescu et al., 2006), due to the lack of Mfd structures in complex with nucleotide, the structural rearrangements of Mfd in the nucleotide catalytic cycle has until now remained unknown. In this study, we provide the first nucleotide bound structure of MsMfd, which to the best of our knowledge represents the first structure of any Mfd in complex with nucleotide. The ADP bound MsMfd complex structure was obtained by co-crystallisation of MsMfd with ATP. After reaction with ATP, the hydrolysis products ADP and inorganic phosphate (Pi) were bound in the crystal structure (Figures 2A and 3D). The two RecA-like domains of the translocation module (TD1/D5 and TD2/D6) of *mycobacterial* Mfd contain seven ATPase signature motifs similar to other SF2 helicase motor proteins (Gorbalenya, Koonin et al., 1989). These motifs are juxtaposed with the nucleotide binding pocket in MsMfd^ADP^ (Figures 1A and 3D). The bound ADP makes van der Waal’s and hydrogen bond interactions with motif I (Walker A), the TD1 loop, and a water mediated hydrogen bond with motif V (Figures 2A and S4A). The released P_i_ is bound to Ser920 in motif V, and in addition makes contacts with Gln709 and Trp900 (Figure 2A).

In this study a SO_4_ molecule occupies the position of the β-Phosphate of the ADP in motif I of the nucleotide binding site in both the selenium derivatized MsMfd and native MsMfd crystal structures. The SO_4_ molecule in both MsMfd^Se^ and native MsMfd structures mimics the effect of β-Phosphate of the ADP substrate in the MsMfd^ADP^ structure (The Root Mean Square Deviation (RMSD) of MsMfd^Se^ and native MsMfd with MsMfd^ADP^ are 0.9 Å and 0.6 Å (across all 1149 and 1168 Cα pairs, Figure S4B)). Hence the SO_4_ molecule drives the active site structural conformation in both MsMfd^Se^ and native MsMfd crystal structures to nucleotide bound state. In the native MtbMfd crystal structure, the nucleotide binding pocket is not occupied by SO_4_ and thus depicts the apo conformation. To gain insights into the structural plasticity of Mfd upon nucleotide binding, we compared the nucleotide bound MsMfd crystal structure with the apo MtbMfd (The RMSD of MsMfd^ADP^ structure with apo-MtbMfd is 2.95 Å (across all 1126 pairs)). Since Mfd is a highly conserved bacterial protein we also compared it with its homologue apo EcMfd crystal structure (PDB-2EYQ, RMSD 8.821 Å (across all 1107 pairs)). MsMfd and MtbMfd share 79% sequence identity at the primary sequence level, and share ∼ 33 % sequence identity with EcMfd. Overall the Mfd structural domain organization is conserved in both *E. coli* and *Mycobacterium* (Figure S2, Figure 1A).

The comparison of TD1/TD2 of MsMfd^ADP^ and apo MtbMfd (Figure 2D) reveals a series of conformational changes. Upon nucleotide binding, the Walker A motif in MsMfd^ADP^ shifts from a compact helical conformation to a looser expanded conformation (Figure 2B and Movie S4). We observe that the pyrimidine ring of ADP is anchored by π-π stacking between the conserved Phe641 in the TD1 loop and a CH-π interaction with Pro822 of the TD1/TD2 hinge-region (Figure 2A). Notably, in apo MtbMfd, and also in apo EcMfd these conserved residues are displaced away from the nucleotide pocket (Figures 2C, 2D and 3B). Thus, the stacking of Pro822 in the hinge region and Phe641 of the TD1 loop against the nucleotide adenine base, strengthens the binding of the nucleotide in MsMfd^ADP^. Unlike ADP bound MsMfd, the hinge regions in apo MtbMfd and apo EcMfd are positioned away from the nucleotide binding pocket and interact with the TRG motif in TD2 (Figures 2C and 2D). It is known that the Mfd TRG motif mediates dsDNA translocation driven by ATP (Chambers, Smith et al., 2003, Mahdi, Briggs et al., 2003). These findings indicate that the nucleotide induced conformational changes in the hinge-region simultaneously strengthens the nucleotide binding, and unlocks the TRG motif through loss of intereactions, leaving the TRG motif of TD2 available for translocation during the ATP catalytic cycle.

### A Flip-flop action of the linker and hinge regions

A flexible linker region connect the MfdN and MfdC domains (Figure S2). Cleavage of the wildtype EcMfd flexible linker enhances its ATPase activity by nearly 20 fold. This may be due to an auto-inhibitory effect arising from MfdN:MfdC domain interactions (Murphy, Gong et al., 2009). To unravel the role of flexible linker and domain interactions in Mfd, we compared the structures of apo and nucleotide bound Mfd. Interestingly, in MsMfd^ADP^ we observe the flexible linker has a marked conformational shift (>8 Å) away from the nucleotide binding pocket compared to apo MtbMfd (Figure 3B). ADP binding and the subsequent movement of the flexible linker lead to the loss of D3:D5 interactions in MsMfd^ADP^ whereas these domain interactions are present in apo MtbMfd (Figure 3A and Movie S2). We observe the presence of D3:D5 interactions even in apo *E. coli* Mfd which drive the flexible linker to adopt a similar conformation to that observed in apo MtbMfd (Figures S5B and S5C; Tables S1A-S1C). The flexible linker and hinge-region swap positions through a flip-flop action that enables the flexible linker to avoid steric occlusion with Phe641 which subsequently stacks against the adenine base of the ADP. Collectively, the nucleotide bound and apo Mfd structures reveal the movements of the flexible linker, the conserved Phe (Phe641 in MsMfd) of the TD1 loop, and the TD1/TD2 hinge-region, which likely function as sensors for the presence or absence of nucleotide. Our structures reveal a set of distinct conformational changes that is directed by the entry and subsequent anchoring of nucleotide at the active site of Mfd. Binding of nucleotide alters the network of non-covalent interactions at the active site (Figure S4A). Although the functionally uncharacterized D3 domain has been proposed to be species-specific with an as yet unknown function (Deaconescu et al., 2006), we observe that the structural fold of the D3 domain is conserved between *E. coli* and *Mycobacterium spp*. As described above, the gain and loss of D3:D5 interactions which influence the flexible linker conformation suggests a previously undiscovered role for the D3 domain in nucleotide binding (Figure 3A; Movies S2 and S3).

### Mfd DNA translocation mechanism and a proposed DNA path

Due to the lack of Mfd structures in complex with nucleotide and/or DNA, the structural mechanism of Mfd translocation is poorly understood. Recently, by using the unzipping mapper technique on EcMfd, Le and co-workers proposed a model for Mfd forward translocation. In this model, nucleotide binding and hydrolysis leads to alternate stepping of TD1 and TD2 on the DNA towards the stalled RNAP (Le et al., 2017). Comparison of the crystal structures of nucleotide bound MsMfd with the apo EcMfd (PDB-2EYQ) revealed that the TD2 of Mfd is rotated outwards by ∼ 46° (Figure 2C). This rotation of TD2 in MsMfd^ADP^ brings the TD2 ATPase motifs and nucleotide-binding site into close proximity (Figure 3D) to participate in the nucleotide catalytic cycle which drives DNA translocation. This mechanism is similar to that observed in other nucleotide bound SF2 helicase structures(Caruthers & McKay, 2002, Sengoku, Nureki et al., 2006, Velankar, Soultanas et al., 1999) (Figures S7A, S7B, S7D and S7E) eg. a rotation of ∼ 101° was observed between TD domains of nucleotide free SsRad54-dsDNA(Durr, Korner et al., 2005) and MsMfd^ADP^ (Figure 3C). Mfd also contains a RecA like translocase domain similar to SF2 *Sulfolobus solfataricus* Rad54 (SsRad54). To further delineate the Mfd DNA translocation pathway, we modelled the MsMfd^ADP^-dsDNA complex using the SsRad54-dsDNA complex crystal structure (Durr et al., 2005, Mazin, Mazina et al., 2010). Based on our MsMfd^ADP^-dsDNA complex model we propose that : (i) the DNA binding mode and directionality observed in the modelled MsMfd^ADP^-dsDNA complex (Figure 3C), is similar to that of the SsRad54-dsDNA complex, (ii) the conserved basic residues in TD1 and motif IV of TD2 which point towards the DNA duplex seem to participate directly in DNA binding (Figure S7F and circled region in Figure 1G).

### MtbMfd rescues the stalled TEC

Electrophoretic Mobility Shift Assay (EMSA) studies show that MtbMfd displaces the RNAP from a stalled transcription elongation complex but not from initiation complexes (Figure 4A), in a similar fashion to EcMfd (Park et al., 2002). The addition of MtbMfd in the reaction mixture leads to the dissociation of the majority of the elongation complex in a concentration and time-dependent manner (Figures 4B and 4C). We observe that while the catalytic activity of MtbMfd is ATP or dATP dependent (Figures S6A and S6B), other nucleotides such as ADP, GTP, CTP and ATPγS do not support activity suggesting that ATP or dATP binding and subsequent hydrolysis is essential for MtbMfd function.

### MtbMfd-RID/RNAP-β subunit interface and specificity

To gain insights in to the specificity of interactions at the interface of mycobacterial Mfd and RNAP, we also modelled the MtbMfd-RID/RNAP-β subunit complex (Figure S6C). Earlier studies demonstrate that the critical residues involved in interactions between the Mfd RNAP Interacting Domain (RID) and the RNAP exist in a bipartite mode, and are phyla specific (Deaconescu et al., 2006, Smith & Savery, 2005, Westblade, Campbell et al., 2010). Our MtbMfd-RID/RNAP-β subunit model reveals that one of these critical residues, Arg540 in MtbMfd-RID, makes a salt-bridge with MtbRNAP Glu132 (Figure S6C and Table S1E). In addition, the highly conserved IKE motif of the β-subunit in bacterial RNAP (Smith & Savery, 2005) is replaced by the IKS motif in MtbRNAP. Our EMSA studies also support the hypothesis that Mfd and RNAP have phyla-specific interactions. This was indicated by displacement activity assays, in which the MtbMfd displacement activity is optimal in the presence of mycobacterial RNAP as opposed to EcRNAP (Figures 4B and 4D).

### EM studies reveal a previously unknown dodecameric assembly of MtbMfd

Size exclusion chromatography (SEC) studies indicate that MtbMfd exists in monomeric and oligomeric hexamer forms (Prabha et al., 2011) in contrast to the exclusively monomeric form of EcMfd (Deaconescu et al., 2006). Intriguingly, negative-stain and cryo-EM solution state maps of the oligomeric SEC fraction of MtbMfd (Figure S1A) revealed a previously undetected dodecameric form (Figures 1E and S3) suggesting that a higher order oligomeric storage form even higher than what was suggested earlier exists. It has been proposed that Mtb may have evolved an oligomeric form of MtbMfd that can be readily released as monomers to participate in TCR as a survival mechanism under stress conditions (Prabha et al., 2011).

## DISCUSSION

Based on apo and nucleotide bound Mfd crystal structures, MsMfd^ADP^-dsDNA model, and in the light of recent biochemical evidence (Le et al., 2017) we present here a comprehensive model for mechanism of Mfd during TCR, that involves domains association and dissociation upon ATP binding, hydrolysis and conformational dynamics during DNA translocation (Figure 4E). In apo Mfd, interactions occur between the D1:D6 (TD2), D2:D7, and D3:D5 (TD1) domains (Figure 4E-i). In TCR, the binding of Mfd to the stalled TEC and subsequent binding of ATP to Mfd might lead to the loss of contacts between the D1:D6 and D3:D5 domains in Mfd (Figure 4E-ii). In this state, the D6 domain rotates freely in such a way that the TRG motif can migrate in a closed conformation towards the upstream minor groove of DNA to initiate translocase activity. This closed conformation can be observed in the 2B subdomain (corresponding to the Mfd TRG motif) of ZfRad54-SO_4_ (Thoma, Czyzewski et al., 2005) (Figure S7C) and is modelled in SsRad54 (Durr et al., 2005). ATP hydrolysis triggers DNA translocation through the TRG motif pushing on the upstream minor groove. Upon ATP hydrolysis the ADP bound Mfd drives the TRG motif to an open conformation, as evidenced in the MsMfd^ADP^ structure (Figures 3C and 3D) and allows Mfd to re-establish D1:D6 interactions (Figures 1D and 4E-iii; Table S1D). At the point of ADP release and ATP binding, the closed conformation positions the TRG motif above the newly translocated upstream DNA site, thus enabling Mfd to move forward during translocation with a ratchet-like motion, where the TRG motif functions as a pawl. Once Mfd reaches the DNA damage site, the D2 domain (UvrA binding domain) of Mfd is untethered from the D7 domain to recruit the UvrA2-B complex (Deaconescu et al., 2006, Deaconescu, Sevostyanova et al., 2012, Fan et al., 2016, Graves et al., 2015, Howan et al., 2012, Selby & Sancar, 1995) (Figure 4E-iv). Subsequently, upon placing the UvrA2-B complex in the immediate vicinity of the DNA damage site the Mfd-RNAP complex is released from DNA in an ATP dependent manner (Fan et al., 2016, Ho, van Oijen et al., 2018) (Figure 4E-v). The UvrA binding determinants of D2 (Manelyte, Kim et al., 2010) are also conserved in the *Mycobacterial* Mfd and are masked by the D7 domain in both apo (not shown) and ADP bound structures (Figure S8).

Structural and functional information on the transcription-coupled DNA repair mechanism in *Mycobacterium,* and in particular from *Mtb*, a human pathogenic bacteria, has until now been poory understood. To our knowledge, this is the first comprehensive structural study on *Mycobacterium* Mfd and will provide a basis for more sophisticated *in vitro* models of Mfd activity in *Mycobacterium*. In summary, the structural snapshots of nucleotide-bound and -free states of the *Mycobacterium* Mfd provide detailed insights into the landscape of the nucleotide binding pocket and conformational remodelling during the nucleotide binding catalytic cycle. The complex and the intricate conformational changes expose the molecular mechanism of Mfd translocation during transcription-coupled DNA repair, which is in all likelyhood generally applicable to all Mfds since Mfds are widely conserved across bacteria. Our functional studies confirm the importance of Mfd in rescuing stalled TEC in *Mycobacterium* species. Furthermore, the structural architecture of oligomeric MtbMfd from solution state cryo-EM shows an unexpected dodecameric assembly, which may represent a mechanism for Mfd storage and release during *Mycobacterium tuberculosis* survival under hostile conditions. Recent studies have shown that Mfd accelerates the development of anti-microbial resistance in pathogenic bacteria including *M. tuberculosis* and is thus an ideal target for anti-evolution drugs (Ragheb et al., 2019). Taken together, these first insights into the three dimensional architecture and structural dynamics during nucleotide-binding in *Mycobacterium* Mfd’s provide a promising basis for developing novel therapeutic agents targeting the Mfd in *Mycobacterium tuberculosis* and other human bacterial pathogens.

## Materials and Methods

### Cloning, expression and purification

The *MsMfd* gene was amplified from *Mycobacterium smegmatis* (*strain mc(2)155*) genomic DNA and cloned into bacterial expression vector pET28a (pETMsMfd) at the NdeI and BamHI restriction sites. The *MtbMfd* gene ((*H37Rv strain*), was isolated and cloned in pET28a (pETMtbMfd) at the restriction sites NdeI and HindIII.

### Overexpression and Purification of *Mycobacterium* Mfd proteins

The overexpression and purification of the MsMfd and MtbMfd proteins were performed using the same protocol unless otherwise specified. pETMsMfd and pETMtbMfd were transformed separately into Rosetta (DE3) (Novagen) competent cells; transformants were selected on LB-Agar plates supplemented with kanamycin (33 µg/ml) and incubated at 37 °C overnight (O/N). The positive transformants were inoculated into 25 ml Luria-Bertani (L.B.) broth as a starter culture and used as a seed for mass culture of 1 litre L.B. broth supplemented with kanamycin (33 µg/ml). IPTG induction was carried out at log phase (0.6 O.D.) at a concentration of 0.8 mM and an induction temperature of 30 °C. The induced culture was grown for 5 hours 30 minutes post IPTG induction, after which the cells were harvested by centrifugation at 8000 rpm for 10 minutes at 4°C. The crude protein lysate was obtained from *E. coli* cells by sonication [in 50 mM Tris (pH 8.0), 500 mM NaCl, 10 % Glycerol, 5 mM Imidazole, 1 mM PMSF, supplemented with EDTA-free protease cocktail inhibitor (Roche)] 15 rounds with 10 sec pulse ON and 30 sec pulse OFF (on ice), followed by centrifugation at 29000 rpm for 1h 45 minutes at 4 °C. The supernatant obtained was again subjected to 55% Ammonium sulfate precipitation. The precipitated mixture was again centrifuged at 13000 rpm for 40 mins at 4 °C. The pellet obtained was then dissolved in solubilization buffer (50 mM Tris (pH 8.0), 500 mM NaCl, 10 % Glycerol, 5 mM Imidazole, and 1 mM PMSF).

The solution was passed through a pre-equilibrated Ni-NTA resin column, and the bound protein was eluted using the elution buffer [50mM Tris (pH 8.0), 500mM NaCl, 10% Glycerol, 10mM β-Mercaptoethanol (βME), and 300mM Imidazole]. Ni-NTA purified protein samples were passed on to heparin sepharose affinity chromatography, and the bound protein was eluted by NaCl gradient (100mM NaCl to 1M NaCl). The samples collected were dialyzed against Equilibration Buffer (EB) (20mM Tris pH 8.0, 500mM NaCl, and 10mM βME) overnight at 4 °C. The resulting protein fractions were concentrated to 1 ml volume using an Amicon centricon (10 kDa MWCO). The concentrated protein was injected into prepacked HiLoad 16/600 Superdex 200 Prep Grade column (GE Healthcare), equilibrated with 2 Column Volume (CV) of EB. The monomer and hexamer fractions were analyzed by 10% SDS-PAGE. The monomer fractions were concentrated to 20 mg/ml and were used for crystallization. For Seleno-methionine labeling (MsMfd**^Se^**), the MsMfd**^Se^** was purified by the feedback inhibition method for experimental phasing(Doublie, 1997). A starter culture was grown overnight in LB media containing kanamycin (33 µg/ml) and was then added into the mass culture and centrifuged. The pellet was dissolved in 10 ml of 1X M9 minimal media and then added into 500 ml of 1X M9 minimal medium which contained kanamycin (33 µg/ml). 15 minutes before IPTG induction, the feedback cocktail inhibitors [Lysine, Threonine, and Phenylalanine(75 mg/L each), Leucine, Isoleucine and Valine (37.5 mg/L each) and L-selenomethionine (45 mg/L)] were added. The culture and pellet were then processed in the same way as for native MsMfd. The pure monomer fractions of MsMfd, MsMfd**^Se^** and MtbMfd samples were stored in 20 mM Tris (pH 8.0), 500 mM NaCl, 10 mM βME and 20 mM Tris (pH 8.0), 200 mM NaCl, 10 mM βME, 2.5% glycerol and 0.25mM EDTA, respectively.

### Crystallisation and cryo preservation of *Mycobacterium* Mfd’s

The MsMfd and MsMfd-ADP protein were crystallised using the hanging drop method at 295 K. The MsMfd-ADP complex was obtained by incubating MsMfd protein with ATP for 1 hour at 4 °C. The reservoir (crystallant) consisted of 100 mM HEPES sodium pH 7.2, 0.2 M Na_2_So_4_, 20% PEG3350 (reservoir and protein were mixed in a 1:1 ratio). Crystals were flash-cooled in a liquid nitrogen using crystallant buffer with 25% PEG3350. The selenium derivative crystals of MsMfd^S**e**^ were crystallised in the same condition as that of native MsMfd. The crystals were cryoprotected with 6% ethylene glycol in the crystallant before being flash cooled in liquid nitrogen. The MtbMfd protein was crystallised using the hanging drop vapour diffusion method at 277 K. The reservoir consisted of 100 mM HEPES Sodium pH 7.0, 800 mM Ammonium formate, 20% PEG3350, 7.5 mM MgCl_2_.6H_2_O (reservoir and protein were mixed in a 1:1 ratio). Crystals were cryoprotected with 6.5% ethylene glycol and flash cooled in a liquid nitrogen.

### Diffraction data collection and processing

The MsMfd and MsMfd-ADP protein crystals diffracted to 2.75Å and 3.5 Å at 100K. Diffraction data for MsMfd were collected on beamline I03 at Diamond Light Source using a 0.9763 Å X-ray wavelength. The MsMfd-ADP diffraction data were collected using a Bruker Micro Star Ultra 2 Copper rotating–anode generator using a MAR345dtb imaging plate, at the In-house School of Biology, Protein Crystallography X-ray facility, IISER Thiruvananthapuram. Multi-wavelength Anomalous Dispersion data were collected for MsMfd^Se^ crystals at 100 K from these crystals on Proxima 2A beamline at Soleil synchrotron to a resolution of 2.99Å. All the MsMfd, MsMfd^se^ MsMfdATP^gammaS^ and MsMfd^ADP^ purified proteins crystallised as a dimers in the asymmetric unit in the orthorhombic space group P2_1_2_1_2_1_ with similar unit cell dimensions (Table 1). MtbMfd single crystal diffraction data were collected on beamline I03 at Diamond Light Source using an X-ray wavelength 0.9763 Å to 3.6 Å resolution at 100 K. In the case of MtbMfd, the protein crystallised as a monomer in the asymmetric unit in the cubic space group P432. The intensity data for the MsMfd^Se^ crystal were indexed, integrated and scaled using XDS(Kabsch, 2010) with the ‘xdsme’ script (https://github.com/legrandp/Xdsme). The intensity data for rest of the MsMfd (Se, ADP, ATPgammaS) and MtbMfd crystals were processed by using *MOSFLM* (Leslie, 2006) and data reduction by *AIMLESS*(Evans, 2006) in the CCP4-7.0.021 program suite (Winn, Ballard et al., 2011).

### Structure determination, model building and refinement of *Mycobacterium* Mfd’s

Initial attempts to solve the structures of MtbMfd and MsMfd by Molecular Replacement (MR) using *E. coli* Mfd (PDB: entry 2EYQ) failed. Therefore, Selenomethionine labelled MsMfd protein was produced and successfully crystallised. Data were collected at the experimentally determined Se absorption edge peak (0.9790 Å) to a resolution of 2.99 Å. The structure was determined with experimental phases obtained by Se-Met SAD phasing using the automated structure solution pipeline - CRANK2(Skubak & Pannu, 2013). 49 of 52 selenomethionines were located in the asymmetric unit using SHELX and used for phase calculation. This permitted automated model building in CRANK2 which proceeded with combined iterative model building, density modification and phase refinement using BUCCANEER, PARROT and REFMAC5 to yield a partial model. The maps were visualized and manual model building carried out using COOT(Emsley, Lohkamp et al., 2010). Initial structural refinement cycles were carried out using REFMAC5(Murshudov, Skubak et al., 2011) and subsequently by *phenix.refine*(Afonine, Mustyakimov et al., 2010) (Table 1, Figure S1F). This structure was then used as a search model to solve the structure of the 2.75 Å MsMfd by MR with MOLREP(Vagin & Teplyakov, 2010). Alternative cycles of model building by COOT(Emsley et al., 2010) and refinement with *phenix .refine* (Afonine et al., 2010) were used to complete the MsMfd structure (Table 1). The final refinement statistics are summarised in Table 1. The structure of MtbMfd, MsMfd^ADP^ and MsMfd^ATPgammaS^ were determined by MR with MOLREP(Vagin & Teplyakov, 2010) using the MsMfd structure as a search model. Initial refinement was done using the jelly body refinement in REFAMC5 (Murshudov et al., 2011) and subsequent iterative cycles of model building and refinement was carried out with COOT and *phenix, refine* (Afonine et al., 2010) (Table 1). Most of the residues in the polypeptide chain were modelled but some disordered regions were encountered for which the electron density was not of high enough quality to model. RMSD were obtained by superposition of pair of structures with 5Å iteration cut off using Chimera (Pettersen, Goddard et al., 2004). Align (Cohen, 1997) and Chimera (Pettersen et al., 2004) were used for the domain angle rotations and RMSD calculations.

### RNA polymerase (RNAP) displacement assay

#### Templates and plasmids

Transcription templates were generated by PCR amplification of selected segments of plasmid pUC18 carrying the sequence of T7A1 (T7 phage early gene) promoter. The transcribed sequence of the T7A1 promoter is as follows.

5’GAATTCAATTTAAAAGAGTA***TTGACT***TAAAGTCTAACCTATAG***GATACT***TACAGCC**A**GAGAGAGGGAGAAGGGAA**T**CGGGATCC… (+27 nt) ….3’

Where the −35 and −10 consensus sequences are set in bold italics, the start or +1 site is shown in bold A, and +20 stalled site is shown in bold T.

### Purification of RNAP and MtbMfd proteins

Overexpression and purification of MtbMfd was carried out as previously described (Prabha et al., 2011). *M. smegmatis* and *M. tuberculosis* RNAP were purified following the methods described earlier(China & Nagaraja, 2010).

### RNAP displacement assays

The assays were carried as previously described(Chambers et al., 2003). Stalled transcription elongation complexes were formed by nucleotide starvation on a 120 bp fragment of plasmid pUC18 containing T7A1 promoter. This fragment encodes a transcript in which at +1 to +19 positions ATP and GTP must be incorporated (sequence is given in the Templates and plasmids section). Transcription elongation complexes stalled at +20 could therefore be formed by using only ATP and GTP in the reaction and excluding UTP and CTP. For the reaction, end labeled template (∼5000 counts/reaction) was incubated with 100 nM of RNAP (MtbRNAP/MsRNAP/EcRNAP) to form an open complex for 15 minutes at 37 °C. These complexes were further incubated with 1mM ATP, GTP each and 50 µg/ml heparin to ensure a single round of transcription initiation for 10 minutes at 37 °C. Stalled complexes were incubated with 4 mM dATP and 250 nM MtbMfd (unless and otherwise indicated) for 30 min at 37 °C. Complexes were analysed by EMSA (4 %) at 4 °C, and radiolabeled bands were detected using phosphor imaging and quantified by image gauze software.

#### Experimental methods specific for Figure 4A

100 nM MsRNAP was incubated with end-labeled T7A1 template to form a complex at 37 °C for 15 min. These complexes treated with either 50 µg/ml heparin or heparin plus 1 mM NTP (ATP and GTP) to form IC and EC respectively at 37 °C for 10 min. Both the complexes were treated with MtbMfd (1 µM) for 30 min at 37 °C and scored by EMSA.

#### Experimental methods specific for Figure 4B,D

100 nM of each bacterial RNAP (**b**, Mtb and **d**, Ec) was incubated with end labeled T7A1 template and the stalled complex was generated by UTP & CTP starvation. The stalled EC was treated with increasing concentrations of purified MtbMfd (0-1 µM) and 4 mM dATP at 37 °C for 30 min. The release of RNAP from template DNA was scored by EMSA.

#### Experimental methods specific for Figure 4C

EC was treated with 250 nM of MtbMfd at 37 °C in a time dependent manner. The rest of the procedures were carried out in the same manner as desribed above.

#### Modelling of the MsMfd^ADP^-DNA and MtbMfd-RID / RNAP-β subunit complexes

The model MsMfd^ADP^-DNA complex (Figure 3C) was generated upon superposition of the ATPase core translocase module of SsRad54-DNA (PDB-1Z63) onto the ATPase core of MsMfd^ADP^ (r.m.s. deviation 2.6 Å for 107/292 Cα pairs), which allowed the double-stranded DNA to be placed into the MsMfd^ADP^ translocase domain junction. The structural complexes of *Thermus thermophilus* (*Tth)* Mfd–RID / *Thermus aquaticus* (*Taq)* RNAP-β1(PDB-3MLQ) and *M. tuberculosis* CarD/RNAP-β subunit (PDB-4KBM), guided the generation of the structural model of MtbMfd-RID/ MtbRNAP-β subunit presented here (Figure S6C). The modelled structures were energy-minimized (with the *Tools - Structure Editing – Minimize structure* option) using the AMBERff14SB force field with 100 steepest descent steps and ten conjugate gradient steps in Chimera to avoid clashes.

#### Negative stain Electron Microscopy sample preparation, data collection and data processing of MtbMfd oligomer

The negative stain sample was prepared by staining the protein with 2% uranyl acetate on a continuous carbon copper grid. The EM images were collected on a JEOL 1200-EX microscope at a voltage of 100 kV at EMBL, Grenoble (France). The spherical aberration of the objective lens was 3.2 mm. 100 digital micrographs were recorded on a 2k x 2k CCD at various defocus values. The images were collected at a magnification of 52974x with a physical pixel size of 15 μm on the CCD (Figure S3A). The image processing was performed using RELION(Scheres, 2012). 2478 particles were manually picked from the 100 micrographs. The CTF estimation was performed using CTFFIND3(Mindell & Grigorieff, 2003). Reference-free 2D classification was performed by grouping 2478 particles into 150 classes. The 2D class average appears to be tetramer-like in structure as shown in Figure S3B. The tetramer-like assembly was different from the earlier finding of a hexameric form(Prabha et al., 2011) of the protein. To rationalize the formation of the tetramer-like assembly, the crystal structure of MtbMfd was analyzed. On generation of the 43 symmetry-related molecules in the P432 space group structure, a tetramer-like assembly (with a tetramer of trimers) was observed. In order to confirm this structure, an initial model was created from the tetramer of trimers in Chimera(Pettersen et al., 2004). The model was a 20 Å map in a box of dimension 190×190×190 pixels and a voxel size of 1.7. For 3D classification, 33 of the 150 2D class averages were considered giving a total of 2048 particles. The reconstruction was performed with an angular sampling of 7.5°. Four 3D class averages were obtained. The class average with the highest signal to noise ratio was used as the initial model for the iterative 3D classification. Further iterative 3D classification with the new model yielded a reconstruction with a resolution of 38.08 Å. This map was again used as an initial model for 3D reconstruction. Repeating the earlier procedure to obtain a refined map, a reconstruction of 30.99 Å was obtained, as shown in Figure S3C. Further repetition of the 3D reconstruction process did not yield any improvement in the resolution of the map. The final map of 30.99 Å resolution was subjected to 3D refinement and local resolution refinement in RELION. ResMap(Kucukelbir, Sigworth et al., 2014) was also used for more precise resolution estimation. The map obtained after ResMap processing has been fitted to the MtbMfd structure with the cubic symmetry applied to generate the dodecamer model and is shown in Figure S3D. A plot of the Fourier Shell Correlation(Scheres & Chen, 2012) (FSC) is shown in Figure S3E. Based on 0.143 criteria for the comparison of two independent datasets, the resolution of the reconstruction is 30.99 Å. The 0.5 FSC line is shown. Thus, while the initial 2D classification appeared to be tetramer-like, on closer observation by using the monomeric purified fraction crystal structure of MtbMfd and knowledge of symmetry, we have modelled the unusual dodecameric state of MtbMfd from the negative stain single-particle electron microscopy imaging and 3D reconstruction.

#### Cryo Electron Microscopy (cryo-EM) specimen preparation, data collection and image processing and three-dimensional reconstruction of MtbMfd oligomer

3 µL of the higher order oligomeric fraction of MtbMfd from SEC (Figure S1A) was applied on holey-carbon Quantifoil R 1.2/1.3 A grids, that had been glow-discharged in air for 90 seconds using a Quorum GloQube unit, adsorbed for 5 s and blotted for 3 to 3.5 s and then plunge frozen into liquid ethane using a vitrobot Mark IV cryo-plunger with its sample chamber maintained at 100 % relative humidity and 18 °C. The vitrified grids were transferred into cartridges. Data was acquired from the grids (maintained at - 186°C) on cartridges over two separate sessions with EPU automated data acquisition software, loaded on an FEI Titan Krios transmission electron microscope operating at 300 kV at InStem-NCBS, Bangalore (India). Each image was collected with an estimated underfocus ranging from 1 to 4 μm on a Falcon III direct detection device (FEI). The dose was fractionated over 79 raw frames collected over a 2-s exposure time (25 ms per frame) with a total dose of 22 e^-^/Å^2^/s. To save space, we stored in 20 frames (bunch of 3 frames together resulting in ∼1.5 e^-^/frame and discarding the last 19). ∼900 movie processed final images were collected, recorded at a nominal magnification of 47,000, corresponding to a pixel size of 1.77 Å at the specimen level. The individual frames were gain corrected, aligned and summed and exposure-filtered according to a dose rate of ∼7 electrons per pixel per second. The image processing was performed using RELION (Scheres, 2012). 397 (out of ∼900 recorded) motion corrected digital micrographs were used for particle picking. With 2D class averages as references from manually picked particles, 48,189 particles were autopicked, sorted and selected to yield a data set of 47,378 particles. The CTF estimation was performed using CTFFIND4 (Rohou & Grigorieff, 2015). Reference-free 2D classification was performed by grouping 16,353 particles into 21 classes. The 2D class average appears to be dodecameric-like in structure as evident from the yellow circled 2D classes in Figure S3F. A preliminary 3D reconstruction was obtained, as shown in Figure S3G. The 3D map was fitted to the MtbMfd structure with the cubic symmetry applied to generate the dodecamer model, as described for the negative stain reconstruction model fitting. Thus, initial 2D classification and 3D reconstruction confirms the existence of a MtbMfd dodecameric state both from the negative stain as well as cryo-EM single-particle electron microscopy imaging and 3D reconstruction.

## Accession codes

All crystal structure coordinates and structure factors have been deposited in the PDB (www.rcsb.org) and EM Maps in EMDB. 6ACA MtbMfd, 6AC8 MsMfd, 6AC6 MsMfd^Se^, 6ACX MsMfd^ADP^ complex. EMD-9614 MtbMfd negative stain reconstruction. MsMfd^ATPgammaS^ complex and cryoEM refined maps can be provided on request.

## ACKNOWLEDGEMENTS

We acknowledge support from the Department of Biotechnology’s (DBT) Ramalingaswamy fellowship grants (to R.N.), and the IISER-TVM fellowship (to S.P.). We thank Soleil synchrotron and CEFIPRA for beam time, funding and local hospitality, in particular A.Thompson and staff of Proxima2A beamline W. Shephard and M. Savko. We thank D. Bellini, P. Lukacik, J. Sandy, J. Weatherby, C. Lobley & K. McAuley at Diamond Light Source. The BM14 diffraction studies at the ESRF (Grenoble) were funded by the BM14 project – a collaboration between DBT (Government of India), EMBL and ESRF. We thank H. Belrahli, B. Manjasetty at EMBL for access to BM14 (ESRF), and K. Manikandan, R. Pillai for their support at EMBL, Grenoble. We thank the Ministry of Human Resource Development (MHRD) IISER Thiruvananthapuram (Government of India) who provided funds for the In-House Protein Crystallography facility at IISER-TVM; Abyson, Eswar and Balachander for their help; the X-ray diffraction facility at MBU, Indian Institute of Science (Funded by DBT and DST, Government of India); NCBS and Mysore University for facilitating crystal screening, in particular M. Vijayan, MRN Murty, N. Suguna, B. Gopal, S. Ramaswamy, NK Loknath, N Vinod and Arif. Cryo-EM data collection was facilitated at the DBT funded National cryo-EM facility in InStem-NCBS, Bangalore. We thank K.R. Vinothkumar and S. Ramaswamy at NCBS and InStem for help with cryo-EM data collection. R.N. thanks in particular Professors E.D. Jemmis and V. Ramakrishnan and his colleagues in the School of Biology. R. N. wishes to dedicated this paper to his father Subramanian Ramanathan who passed away on 1 July 2019.

## AUTHOR CONTRIBUTIONS

S.P. performed cloning, protein expression, purification and crystallization of Mfd’s with the help of R.N. X-ray data collection and crystallographic analysis was performed by S.P., G.A.F. and R.N with help from M.A.W. Sw.P. performed EMSA and RNAP displacement assays, under the supervision of V.N. and D.N.R. who also helped in data analysis. S.P. purified the protein for negative stain and cryo-EM studies. R.N. performed negative stain specimen preparation and Data collection with help of K Manikandan and V.B. along with R.N. performed image processing and 3D reconstruction. R.N. did the cryo-EM image processing and 3D reconstruction. R.N., V.N. and D.N.R. designed and supervised the project. S.P. and R.N wrote the manuscript with input from the other authors.

## Appendix

**Table S1.**
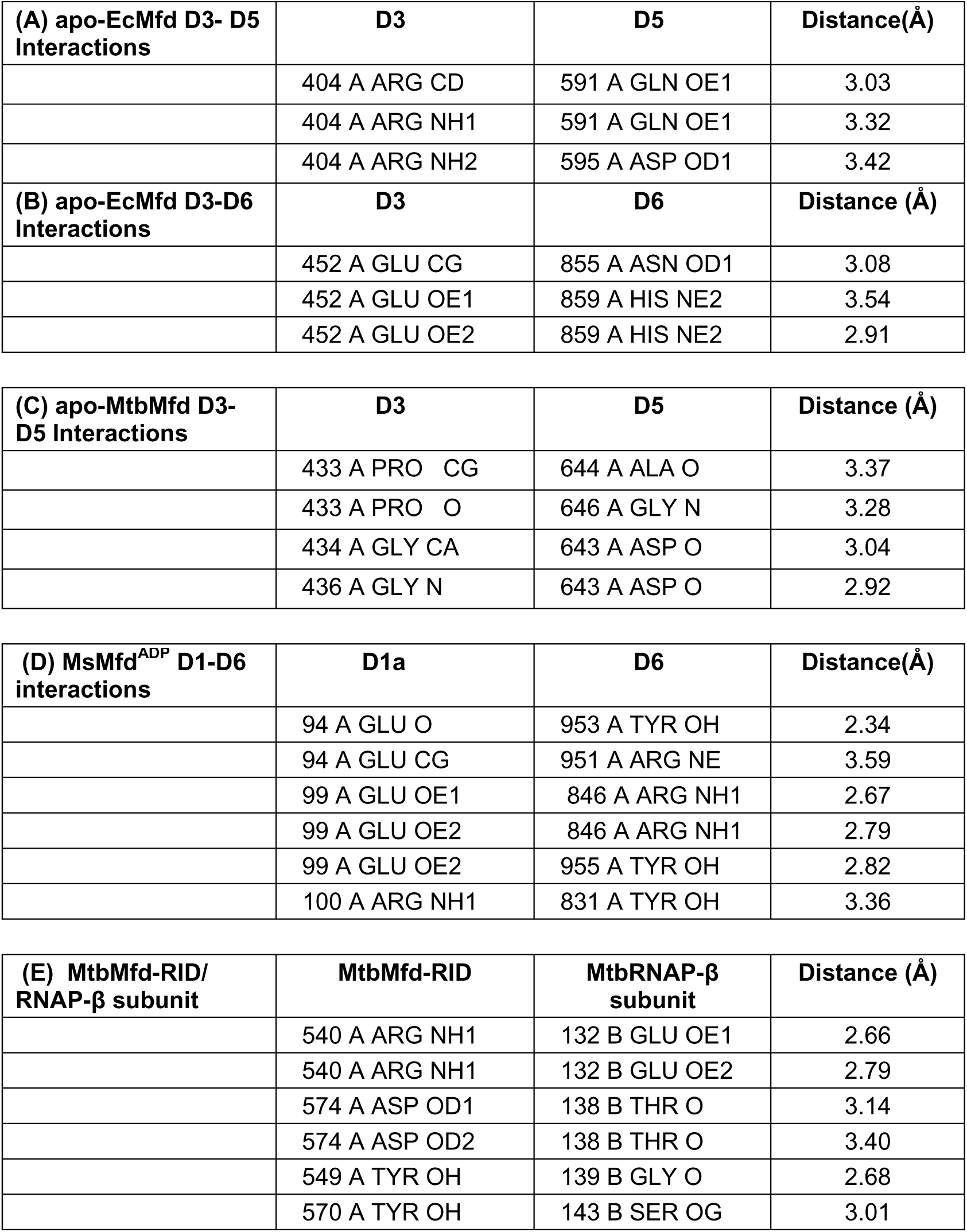
Inter domain interactions from crystal structures of Mfd’s. **(A,C)** D3 domain interaction with the D5 domain in both apo-EcMfd and apo-MtbMfd. **(B)** In addition D3:D6 interactions exist in apo-EcMfd. **(D)** Interactions between the D1a and D6 domains of MsMfd^ADP^. **(E)** The interface contacts in the modelled MtbMfdRID and RNAP-β subunit.

| **(A) apo-EcMfd D3- D5 Interactions** | **D3** | **D5** | **Distance(Å)** |
| --- | --- | --- | --- |
|  | 404 A ARG CD | 591 A GLN OE1 | 3.03 |
|  | 404 A ARG NH1 | 591 A GLN OE1 | 3.32 |
|  | 404 A ARG NH2 | 595 A ASP OD1 | 3.42 |
| **(B) apo-EcMfd D3-D6 Interactions** | **D3** | **D6** | **Distance (Å)** |
|  | 452 A GLU CG | 855 A ASN OD1 | 3.08 |
|  | 452 A GLU OE1 | 859 A HIS NE2 | 3.54 |
|  | 452 A GLU OE2 | 859 A HIS NE2 | 2.91 |
| **(C) apo-MtbMfd D3-D5 Interactions** | **D3** | **D5** | **Distance (Å)** |
|  | 433 A PRO CG | 644 A ALA O | 3.37 |
|  | 433 A PRO O | 646 A GLY N | 3.28 |
|  | 434 A GLY CA | 643 A ASP O | 3.04 |
|  | 436 A GLY N | 643 A ASP O | 2.92 |
| **(D) MsMfd<sup>ADP</sup> D1-D6 interactions** | **D1a** | **D6** | **Distance(Å)** |
|  | 94 A GLU O | 953 A TYR OH | 2.34 |
|  | 94 A GLU CG | 951 A ARG NE | 3.59 |
|  | 99 A GLU OE1 | 846 A ARG NH1 | 2.67 |
|  | 99 A GLU OE2 | 846 A ARG NH1 | 2.79 |
|  | 99 A GLU OE2 | 955 A TYR OH | 2.82 |
|  | 100 A ARG NH1 | 831 A TYR OH | 3.36 |
| **(E) MtbMfd-RID/ RNAP-β subunit** | **MtbMfd-RID** | **MtbRNAP-β subunit** | **Distance (Å)** |
|  | 540 A ARG NH1 | 132 B GLU OE1 | 2.66 |
|  | 540 A ARG NH1 | 132 B GLU OE2 | 2.79 |
|  | 574 A ASP OD1 | 138 B THR O | 3.14 |
|  | 574 A ASP OD2 | 138 B THR O | 3.40 |
|  | 549 A TYR OH | 139 B GLY O | 2.68 |
|  | 570 A TYR OH | 143 B SER OG | 3.01 |

**Figure S1.**
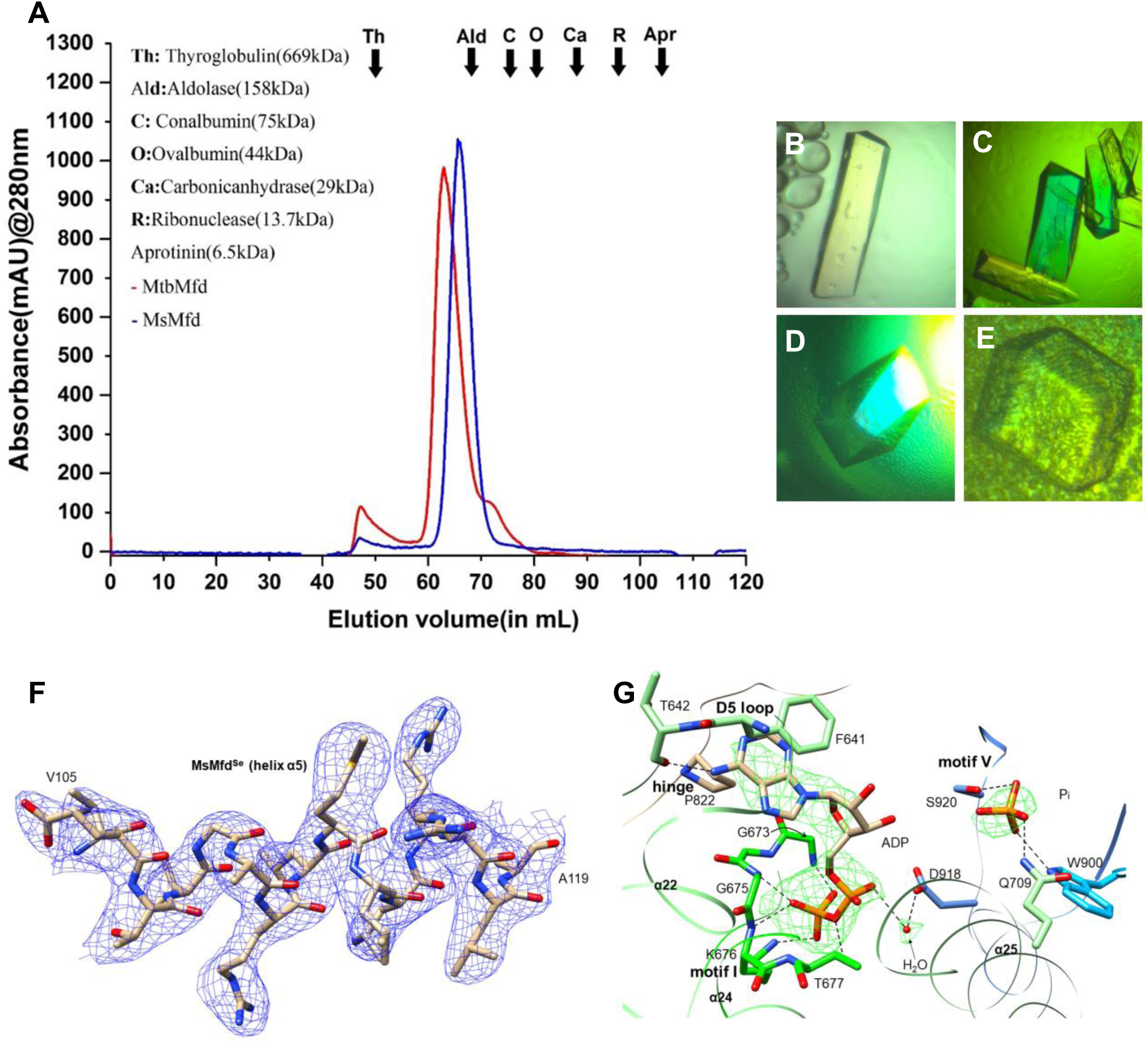
***Mycobacterium* Mfds protein purification and crystallographic data.** (**A**) Size Exclusion Chromatography (HiLoad 16/600 Superdex 200 Prep Grade) elution profile of wild-type MsMfd (blue) and MtbMfd (red) showing hexamer and monomer species. The higher oligomeric form observed for the MtbMfd is consistent with the earlier studies(Prabha, Rao et al., 2011). In this current study we observe that MsMfd elutes predominatly as monomer with a negligible amount of hexamer. The black arrows indicate the peak position of standard proteins. **(B-E)** *Mycobacterium* Mfd crystals. (**B**), (**C), (D)** and **(E)** show crystals of MsMfd, MsMfd^se^, MsMfd^ADP^ (Space Group P2_1_2_1_2_1_) and MtbMfd (Space Group P432). (**F**) Representative quality of the MsMfd^se^ electron density map. 2m*F*_o_ − D*F*_c_ electron density map (contoured at 1.0 *σ)* for an α-helix (α5) in the D1a domain of MsMfd^Se^ refined to 2.99 Å resolution. (**G**) MsMfd^ADP^ *F*_o_ − *F*_c_ map is shown as green mesh for bound ADP, P_i_ and water at the 3σ counter level. The map was generated before modelling of the ADP, Pi and Waters in the electron density.

**Figure S2.**
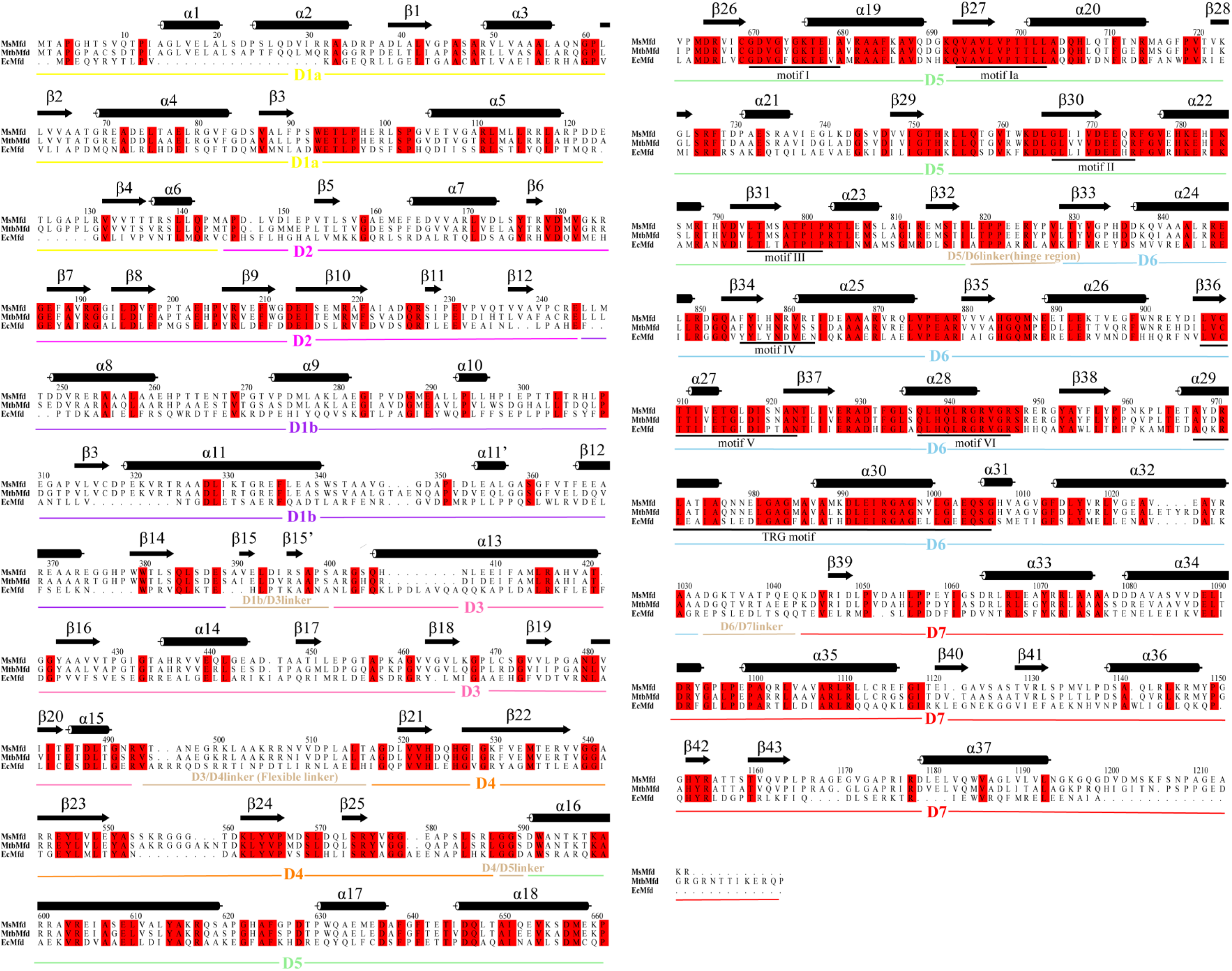
**Multiple sequence alignment of *Mycobacterium* Mfds and *E. coli* Mfd.** The sequence alignment includes *M. smegmatis*, *M. tuberculosis,* and *E.coli* Mfd. The MsMfd sequence is shown at the top position along with its secondary structural elements (obtained from the crystal structure with DSSP(Kabsch & Sander, 1983)). The domain organization for MsMfd is denoted at the bottom of Multiple Sequence Alignment by the colored bars (color-coded as in Figure 1A). The ATPase motifs I to VI and the TRG motifs are indicated with appropriately labelled black bar and the invariant residues are colored in red. The sequence alignment was carried out using Muscle(Edgar, 2004). Rendering of aligned sequences was carried out using Alscript(Barton, 1993). Pairwise alignment of the proteins was carried out with EMBOSS: https://www.ebi.ac.uk/emboss – Needle.

**Figure S3.**
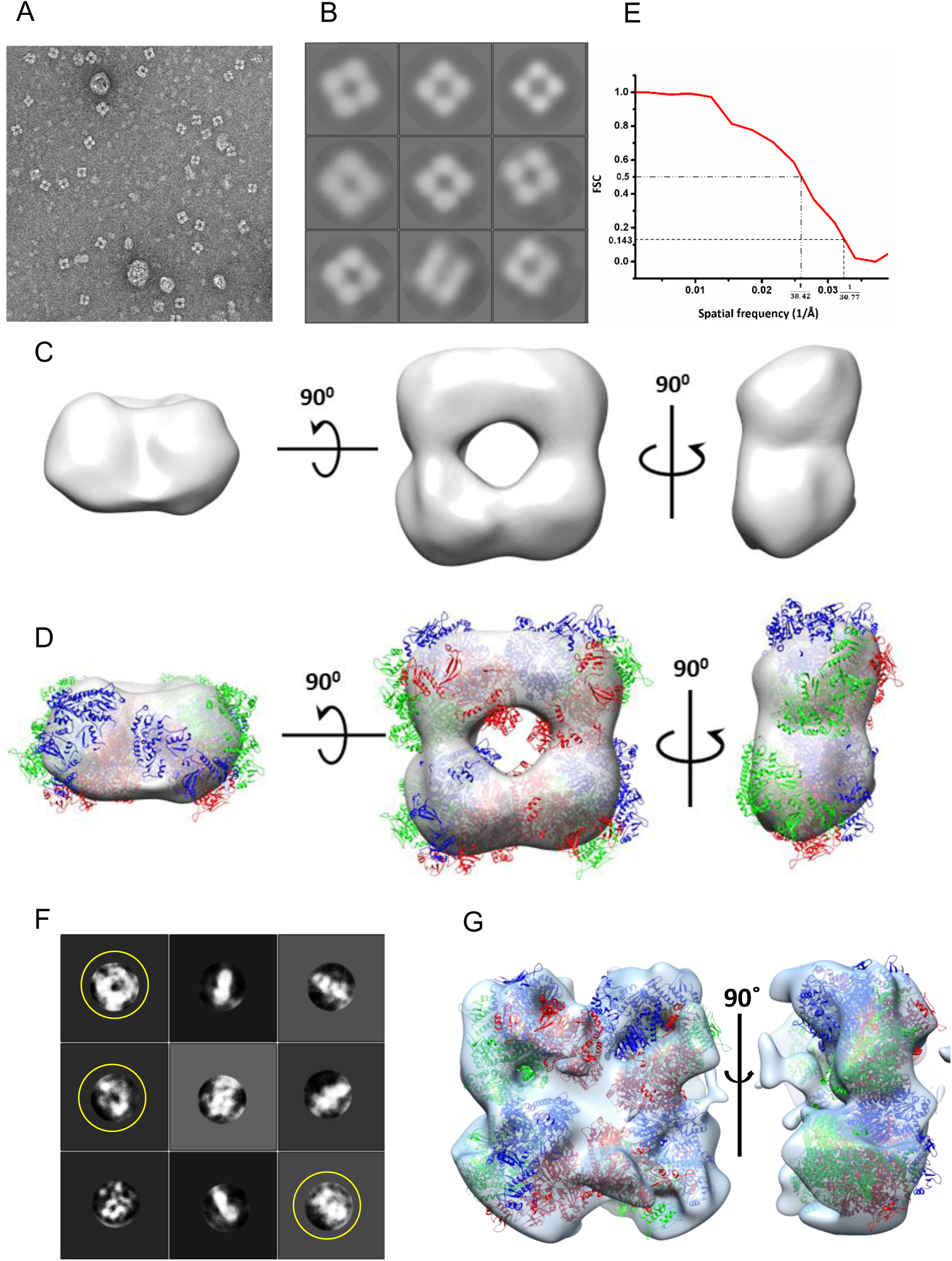
**Negative stain and cryo-EM reconstruction shows a dodecameric form of MtbMfd.** (**A**) MtbMfd oligomer representative digital micrograph clearly showing 4-fold symmetry of domains (**B**) Representative 2D class averages from 150 classes of the 2478 particles that were manually picked from 100 digital micrographs. (**C**) Negative stain EM 3D reconstruction of MsMfd showing a 30.99 Å map contoured at 2 sigma. (**D**) Illustrates the Mfd dodecameric form with fitted MtbMfd coordinates (using MtbMfd asymmetric monomer crystal structure, the 43 - cubic symmetry was applied to obtain the dodecameric form of MtbMfd). This dodecameric form was rigid body fitted to the reconstruction map shown in **c** using Chimera. Each MtbMfd trimer is shown in red, blue and green. (**E**) The Fourier Shell Correlation(Scheres & Chen, 2012) (FSC) plot for 3D reconstruction. (**F**) Representative 2D class averages from cryo-EM data set. The similarity of cryo-EM 2D classes (yellow circled) with the corresponding look alike 2D class projections of the dodecamer assembly from negative stain indicates the existence of similar dodecameric forms in solution state. (**G**) The cryo-EM preliminary C1 asymmetric 3D reconstruction map shows the dodecamer assembly, which is consistent with the negative stain reconstruction observed in Figure S3D. The dodecameric form of MtbMfd coordinates are shown fitted into the cryo-EM map (using Chimera).

**Figure S4.**
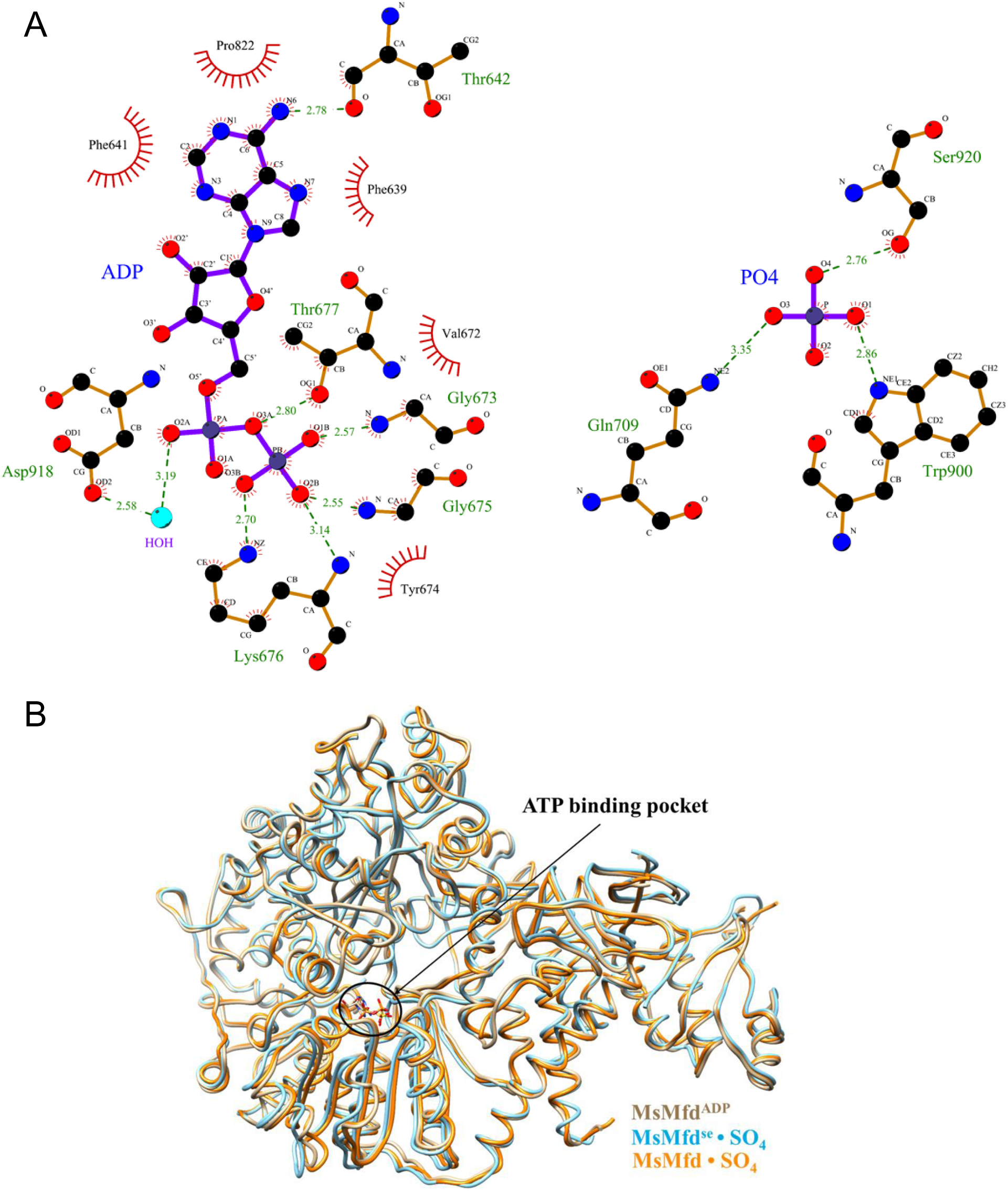
**(A) Schematic diagram of MsMfd-ADP+ Pi complex interactions generated by LigPlot.** Residues which interact with the ADP (left) panel and Inorganic Phosphate Pi (right) in the MsMfd^ADP^ crystal structure are highlighted. The hydrogen bonds (green dashed lines) and van der Waal’s/hydrophobic interactions (red) are shown. (**B**) Superposition of SO_4_^2-^ bound MsMfd structures: MsMfd^Se^ and MsMfd on to MsMfd^ADP^ crystal structure. The SO_4_ is seen bound in the position of the ADP β-phosphate at the ATP binding pocket (circled region). Hence, the native MsMfd (orange) structures bound to SO_4_^2-^ show a structural conformation similar to the nucleotide-bound MsMfd^ADP^ (see main text).

**Figure S5.**
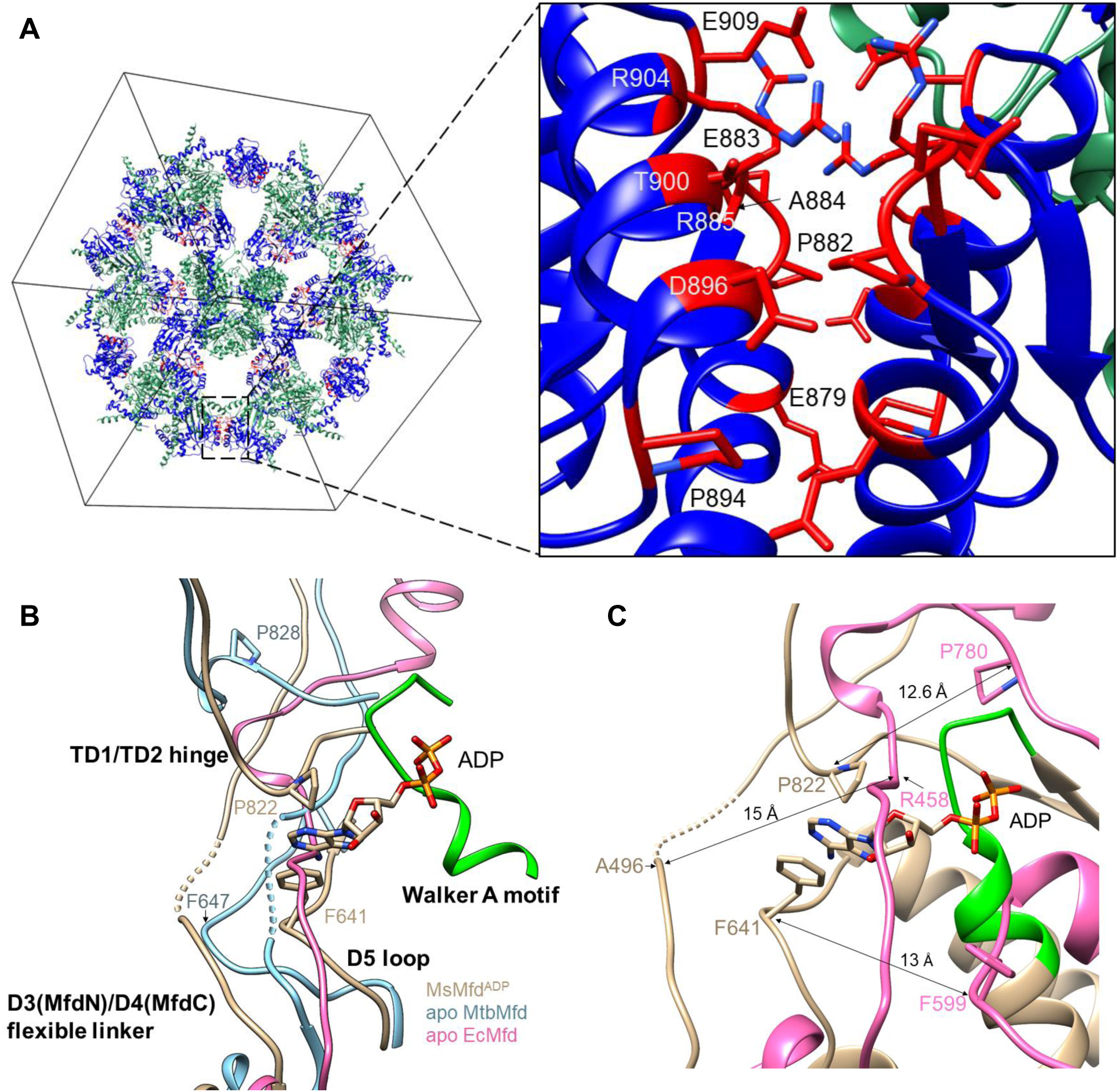
**(A)** The translocation module of apo-MtbMfd in the P432 crystal unit cell. Only TD1 (sea green) and TD2 (blue) are shown with crystal contacts. (Left) The display of crystal symmetries of TD1 and TD2 in the unit cell. Only TD1 and TD2 of the MtbMfd crystal structure is shown for simplicity. (Right, inset) Magnified view of the boxed region showing the residues in the TD2 domain that are involved in crystal contacts (in red). These residues were identified by CONTACT in the CCP4 program suite (Winn, Ballard et al., 2011). **(B,C)** Flip-flop movement of the flexible linker and hinge region in the ATP catalytic cycle: (**B**) Conformational variations of the flexible linker in apo and nucleotide bound Mfds. Upon superposition of apo-EcMfdN on to the MsMfd^ADP^ the flexible linker is positioned near the nucleotide, as seen in apo-MtbMfd. In the nucleotide-bound MsMfd^ADP^ structure, the position of the flexible linker is displaced away from the nucleotide, the Walker A motif is shown in Green. Similar dynamic features observed in both apo-EcMfd and apo-MtbMfd suggest a critical role for the flexible linker, the hinge region and the D5 loop in the nucleotide induced conformational changes. Dotted lines in both figures indicate regions not observed in the electron density. (**C**) Overall superposition of MsMfd^ADP^ (tan) and apo-EcMfd (hot pink) shows loop movements at the nucleotide bound region in a similar manner as seen in MsMfd^ADP^ and apo-MtbMfd, Figure 3B. In apo-EcMfd, the hinge region, D5 loop and flexible linker are displaced by 12.6 Å, 13 Å and 15 Å, respectively with respect to MsMfd^ADP^.

**Figure S6.**
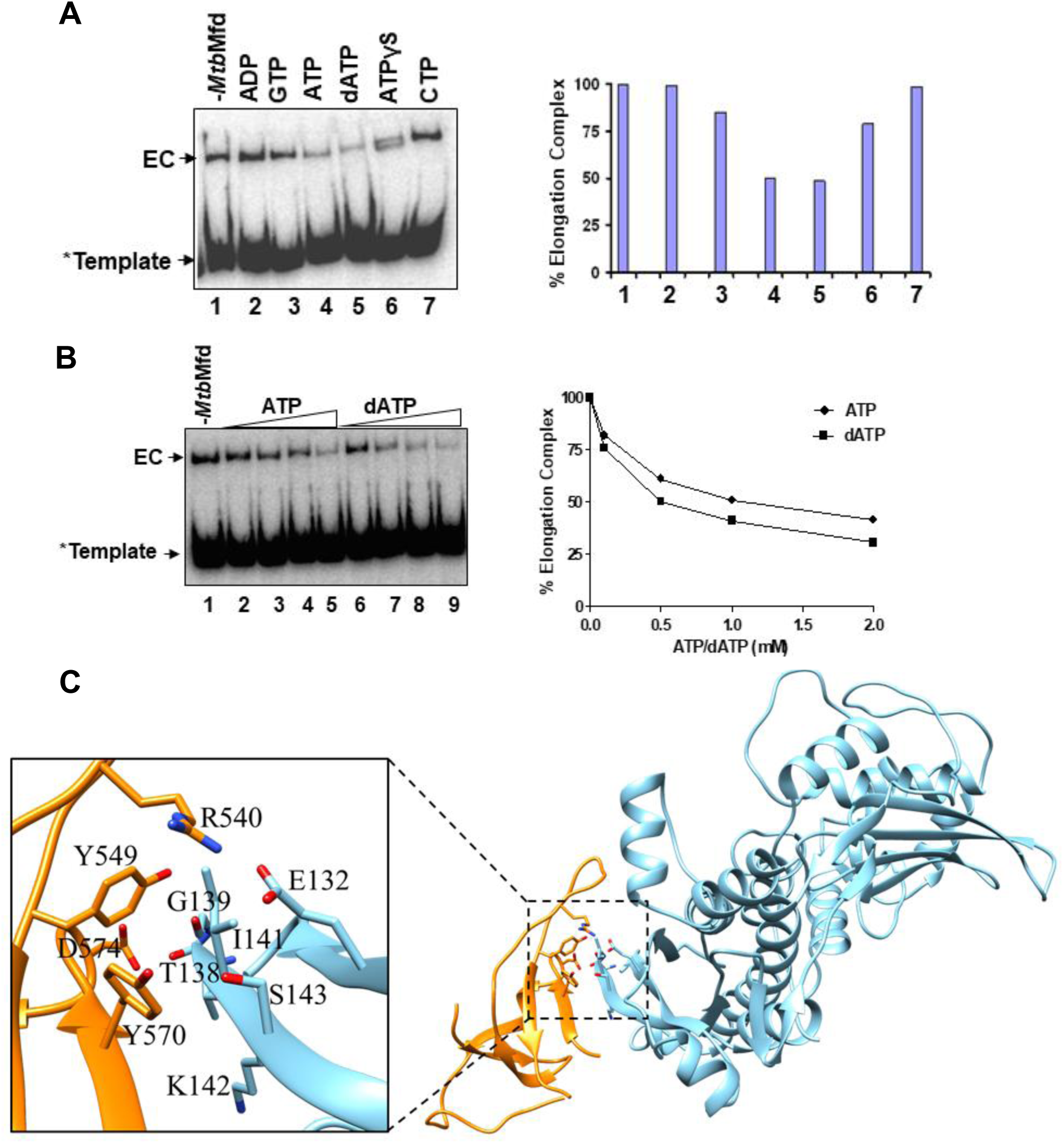
**Effect of nucleotides on RNAP displacement activity of MtbMfd: (A)** Release of RNAP by MtbMfd from the stalled elongation complex (EC) in the presence of different nucleotides (lanes 2-7). Lane 1 represents the elongation complex only in the absence of MtbMfd and nucleotide. The nucleotides (2 mM each) used are highlighted at the top of the gel. Quantification of data obtained is shown in the panel on the right. (**B**) Displacement of RNAP by MtbMfd in the presence of increasing concentrations (0-2 mM) of ATP (lanes 2-5) or dATP (lanes 6-9). Quantification of data obtained is shown in the panel on the right. In both **a** and **b**, the release of RNAP from template DNA was scored by EMSA (4 %) at 4 °C and visualized by a phosphor imager. The EC formed in the absence of MtbMfd is considered as 100%. (**C**) Modelled MtbMfdRID (orange) interaction with the β subunit of MtbRNAP (sky blue). The model was generated as described in the methods section above. The interface of the MtbMfdRID and β subunit of RNAP residues contacts are shown in Table S2E.

**Figure S7.**
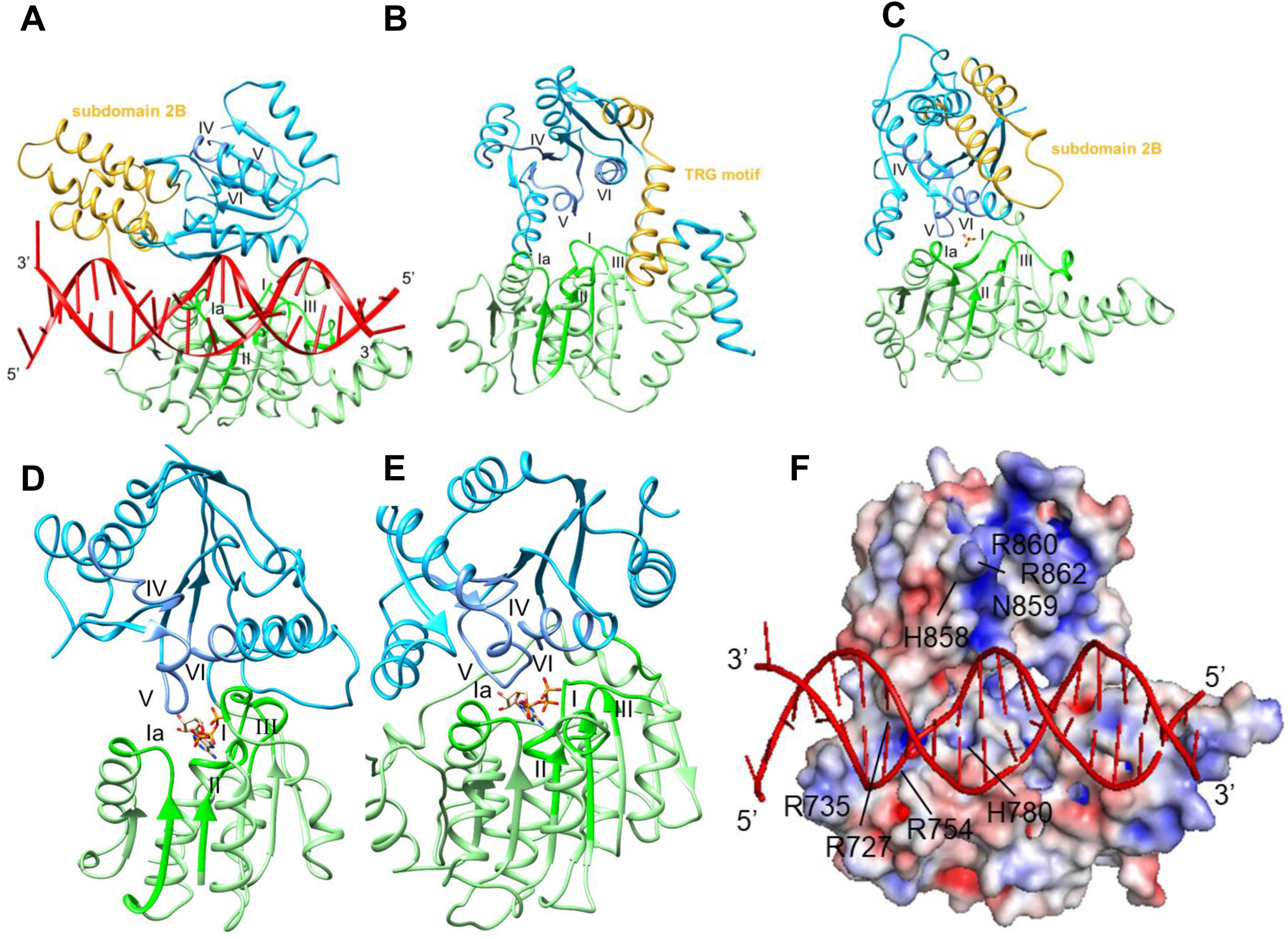
**The RecA-like translocation domains.** (**A**) Nucleotide free SsRad54-dsDNA (PDB1Z63). (**B**) Nucleotide-free EcMfd (PDB2EYQ). (**C**) ZfRad54-SO4 (PDB1Z3I). (**D**) PcrA-ATP-SsDNA (PDB-3PJR). (**E**) Vasa-ANP-RNA (PDB-2DB3). The ATPase motifs of TD1(light green) and TD2 (deep sky blue) domains are shown in green and corn flower blue. **(A, B, C)** The TRG motif of Mfd in (**B)** and its corresponding motif annotated as subdomain 2B in (**A)** SsRad54 and **(C)** ZfRad54 are shown in gold. **(D,E)** The translocation domain (TD2) of PcrA is rotated 21° while Vasa by 15° with respect to TD2 of MsMfd^ADP^ (not shown). Translocase domains (the TD1/TD2 of MtbMfd shows an r.m.s.deviation 1.6Å for 437 Cα positions with that of MsMfd^ADP^; across all 442 pairs:1.8Å). Notably, significant structural differences were observed between the TD1/TD2 of apo-EcMfd (nucleotide-free) and MsMfd^ADP^ TD1/TD2 (RMSD between 300 Cα positions is 2.1 Å; across all 442 pairs: 8.4Å). The ATPase motifs of the translocase domains are positioned closer to the nucleotide-binding pocket in the nucleotide-bound structure (D,E) as compared to nucleotide free structures (A, B). (**F**) Electrostatic surface potential of the translocation module of MsMfd^ADP-^ DNA complex model. The basic residues R735, R727, R754, H780 in TD1 and H858, R860, R862 in motif IV of TD2 along with polar N859 are residues that might interact with DNA during the Mfd translocation event are highlighted. The MsMfd^ADP^ translocation module - DNA complex model was generated by superposition of SsRad54-DNA (PDB-1Z63) on to the MsMfd translocation domains (see the methods section above). The electrostatic surface potentials were calculated with the APBS plug-in in pymol (www.pymol.org) and are contoured from −5(red) to +5 (blue) kT/q.

**Figure S8.**
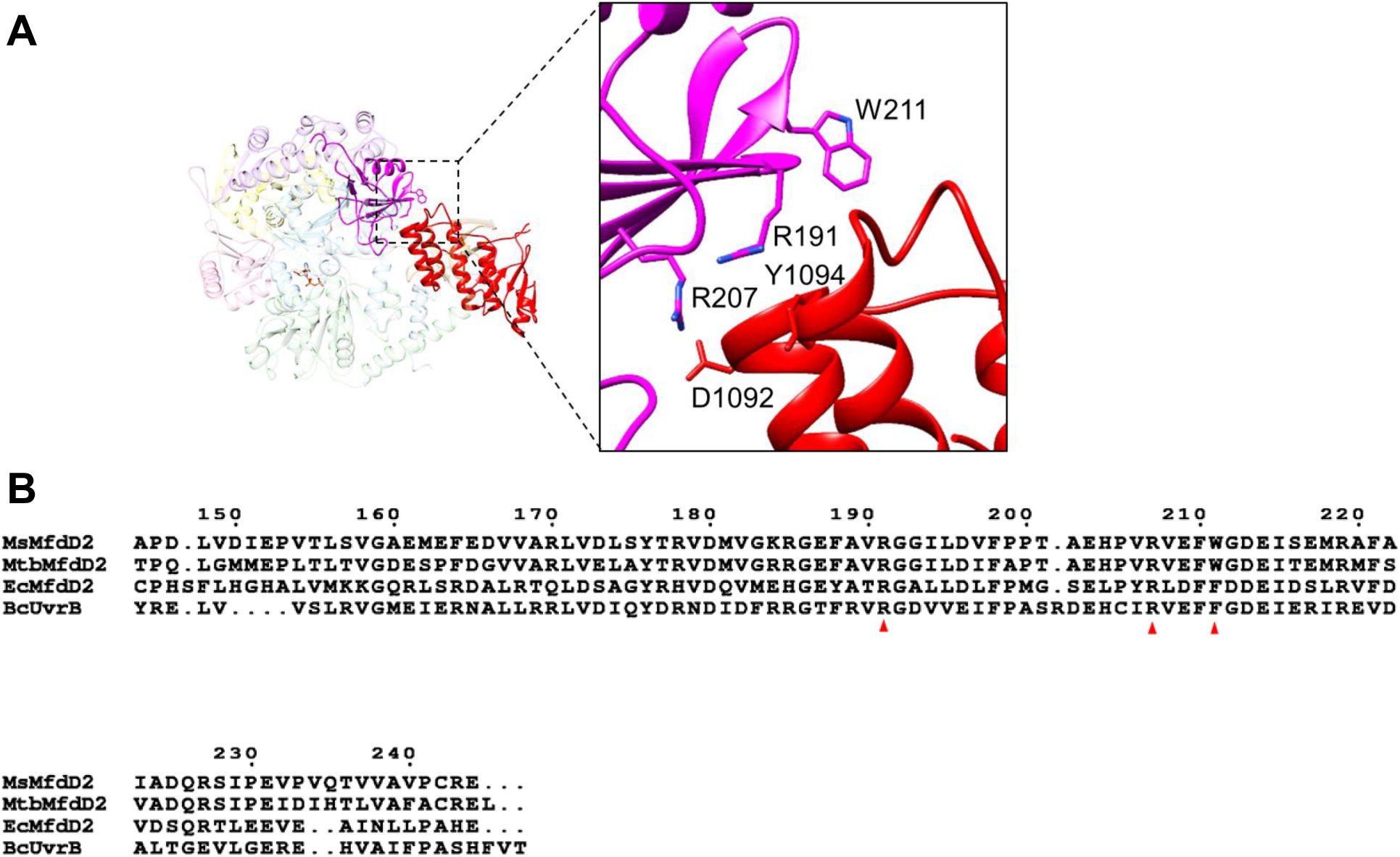
**The D2 domain masked by the D7 domain in MsMfd^ADP^:** (**A**) The UvrA interacting residues (R191 and R207) in the D2 domain (magenta) which interact with Y1094 and D1092 in the D7 domain (red) at the D2/D7 interface are buried and hence are not available for recruiting UvrA (residues relative accessibility was calculated using NACCESS (Hubbard & Thornton, 1993)). The other UvrA interacting residue W211 is seen exposed to the solvent in the vicinity of the D2 and D7 interface. (**B**) Sequence alignment of the D2 of MsMfd, MtbMfd, EcMfd, and BcUvrB. The conserved UvrA interacting residues are highlighted with a red triangle at the bottom of the sequence.

## Appendix Movies

**Movie S1.** Mfd translocation domain TD2 dynamics with respect to TD1 upon nucleotide binding.

**Movie S2.** Nucleotide induced remodelling in mycobacterium Mfds.

**Movie S3.** Nucleotide induced conformational dynamics of MsMfd in comparison with EcMfd.

**Movie S4.** Conformational switching of Walker A motif in presence and absence of nucleotide.

